# Genetic analyses identify widespread sex-differential participation bias

**DOI:** 10.1101/2020.03.22.001453

**Authors:** Nicola Pirastu, Mattia Cordioli, Priyanka Nandakumar, Gianmarco Mignogna, Abdel Abdellaoui, Benjamin Hollis, Masahiro Kanai, Veera M. Rajagopal, Pietro Della Briotta Parolo, Nikolas Baya, Caitlin Carey, Juha Karjalainen, Thomas D. Als, Matthijs D. Van der Zee, Felix R. Day, Ken K. Ong, Finngen Study, 23andMe Research Team, iPSYCH Consortium, Takayuki Morisaki, Eco de Geus, Rino Bellocco, Yukinori Okada, Anders D. Børglum, Peter Joshi, Adam Auton, David Hinds, Benjamin M. Neale, Raymond K. Walters, Michel G. Nivard, John R.B. Perry, Andrea Ganna

**Author notes:** These authors contributed equally to this work (Nicola Pirastu, Mattia Cordioli, Michel G. Nivard, John R.B. Perry, Andrea Ganna).

## Abstract

Genetic association results are often interpreted with the assumption that study participation does not affect downstream analyses. Understanding the genetic basis of this participation bias is challenging as it requires the genotypes of unseen individuals. However, we demonstrate that it is possible to estimate comparative biases by performing GWAS contrasting one subgroup versus another. For example, we show that sex exhibits autosomal heritability in the presence of sex-differential participation bias. By performing a GWAS of sex in ~3.3 million males and females, we identify over 158 autosomal loci significantly associated with sex and highlight complex traits underpinning differences in study participation between sexes. For example, the body mass index (BMI) increasing allele at the *FTO* locus was observed at higher frequency in males compared to females (OR 1.02 [1.02-1.03], P=4.4×10^−36^). Finally, we demonstrate how these biases can potentially lead to incorrect inferences in downstream analyses and propose a conceptual framework for addressing such biases. Our findings highlight a new challenge that genetic studies may face as sample sizes continue to grow.

## Introduction

Individuals who enroll in research studies or who purchase direct-to-consumer genetic tests are often non-representative of the general population^1,2,3^. For example, the UK Biobank study invited ~9 million individuals and achieved an overall participation rate of 5.45%^4^. Enrolled individuals demonstrate an obvious ‘healthy volunteer bias’, with lower rates of obesity, smoking and self-reported health conditions than the population sampling frame^4^. Achieving good representation of the sampled population in any study is a difficult challenge. Some examples do exist, such as the iPSYCH study which gathered a random population sample by extracting DNA from a nationwide routine collection of neonatal dried blood spots and linkage to national register data^5^. The benefits of good representation have been long debated^6,7,8,9^. Many researchers argue that non-representative studies can bias prevalence estimates but do not lead to substantial bias in exposure-disease associations^10,11^. Deliberately non-representative study designs can also be valuable, for example by enriching for cases who carry more disease causing alleles in a case-control study to maximize the power to detect genetic effects^12^.

There is recent evidence that genetic factors are associated with degree of study engagement^13,14,15^. For example, within a study, individuals with high genetic risk for schizophrenia are less likely to complete health questionnaires, attend clinical assessments and continue to actively participate in follow-up than those with lower genetic risk^13,16^. It remains unclear to what extent genetic factors influence initial study enrollment, or what are the downstream consequences of such bias, although previous simulations have attempted to quantify this bias ^17^. We hypothesised that study participation bias can be identified by performing a GWAS on a non-heritable trait. Given that there are no known biological mechanisms that can give rise to autosomal allele frequency differences between sexes at conception, any allele frequency difference between sexes highlights an impact of that locus on sex-differential survival or sex-differential study participation. Another way to describe this concept is, if any trait leads males and females to differentially participate in a study, we would observe artefactual associations between variants associated with that trait and sex (see **Box 1 and Extended Figure 1**). Therefore, an autosomal GWAS of sex provides a unique negative control analysis for genetic association testing, and may provide novel insights into the factors that underlie non-representative study participation^18^.

### Box 1

#### Participation bias

Participation - also called “selection” or “sampling” - bias is observed when participation in a study is not random^38,39^ with respect to the reference population. Participation bias can impact prevalence estimates and results in biased association estimates. This latter phenomenon is caused because participation bias acts as a “collider”.

#### Collider bias

If two variables independently cause a third variable (the collider), then conditioning on the collider (i.e. conditioning on study participation) can cause a spurious association between the two variables^40^. In **Extended Figure 1** we draw 3 path diagrams representing different types of participation bias.

#### Sex-differential participation bias

Sex-differential participation bias is a special case of participation bias where the determinants of study participation affect women and men to differing extents. While participation bias can be detected only if information on non-participating individuals are available, sex-differential participation bias can be detected by comparing genetic allele frequencies between males and females within a study.

Here, we report the results from such a GWAS of sex, performed in ~3.3 million genotyped individuals. We identify more than 150 independent autosomal signals significantly associated with sex, highlighting several complex traits that contribute to sex-differential study participation. Furthermore, we demonstrate the potential impact of such bias on association testing and discuss a conceptual framework to address this issue.

## Results

We performed a GWAS of sex (females coded as 1, males coded as 0) in 2,462,132 research participants from 23andMe using standard quality control procedures (**Supplementary Note**). We identified 158 independent genome-wide significant (P<5×10^−8^) autosomal signals, indicating genetic variants that show significant allele frequency differences between sexes in this sample (**Figure 1** and **Supplementary Table 1**).

**Figure 1:**
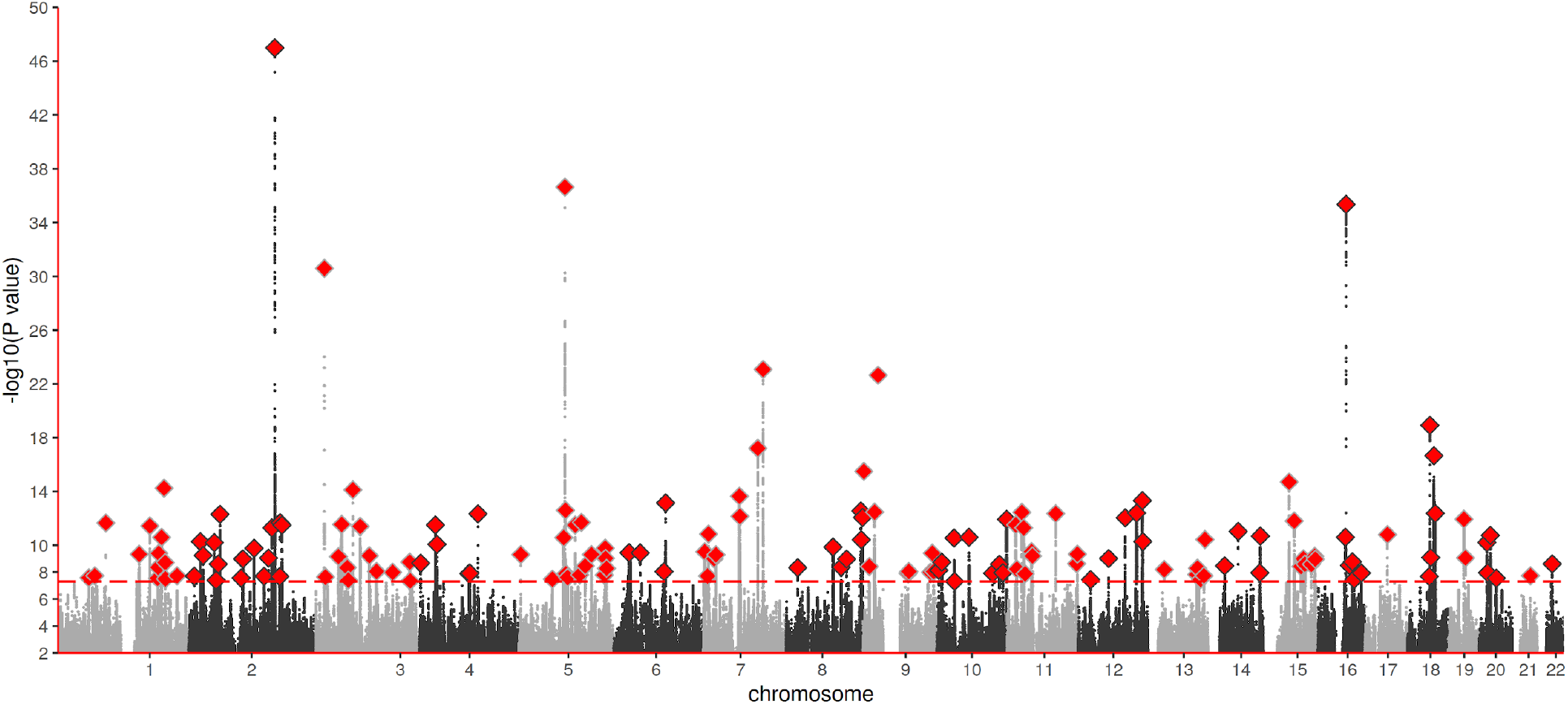
Manhattan plot for a GWAS of sex in 2,462,132 participants from 23andMe.The plot reports all identified loci including those filtered by the extremely stringent quality control applied to directly-genotyped SNPs.

### Technical artefacts do not explain autosomal associations with sex

Additional conservative quality control procedures were performed to exclude any significant signals that might be caused by technical error (**Supplementary Note**). The most obvious reason for a false-positive association with sex is that the autosomal genotype array probe cross-hybridizes with a sex chromosome sequence. This issue has impacted similar previously published studies. For example, a GWAS in 8,842 South Korean males and females identified nine genetic variants strongly associated with sex^19^. The authors attributed their findings to biological mechanisms determining sex-selection, however, all of those nine associated variants are located within autosomal regions with significant homology to a sex chromosome sequence.

To evaluate the impact of this issue in our own data, we first identified directly genotyped variants that were both genome-wide significantly associated with sex and in LD (r^2^>0.1) with one of our imputed top signals (N=78; **Supplementary Table 2**). We then tested for sex chromosome homology with the genomic sequence (+/− 50bp) surrounding each genotyped variant, and found that one quarter (18/78) of our signals were potentially attributable to this technical issue. We further excluded additional loci due to low allele frequency, significant departure from Hardy-Weinberg equilibrium and/or low genotyping success rate, 49/78 directly-genotyped genome-wide significant signals remained. These data suggest that the majority of signals that we identify represent true allele frequency differences between the sampled male and female participants in 23andMe, rather than due to genotyping errors.

### Survival bias does not explain autosomal associations with sex

We next explored whether the observed signals for sex might arise due to sex-differential effects on mortality. To evaluate this, we repeated the GWAS of sex but restricted the sample to individuals aged 30 years or younger (N=320,487), under the assumption that effects due to sex-differential mortality are less likely in younger than older age groups. While the substantially smaller sample size weakened the statistical significance of the signals, the magnitudes of effect across most signals remained consistent (**Extended Figure 2**), with no significant difference in effect size observed for any of the 158 loci (**Supplementary Table 3)**.

### Participation bias results in autosomal associations with sex

We next explored the hypothesis that many signals for sex that act by influencing sex-differential study participation rates may show markedly different associations with sex by study recruitment design (whereas effects due to sex-differential mortality would be consistent between studies). We therefore repeated the GWAS of sex in 4 additional studies - UK Biobank, Finngen, Biobank Japan and iPSYCH (total N = 847,266) - which varied by study recruitment design. As in 23andMe, UK Biobank required active participant engagement, albeit following a very different sampling and recruitment process. By contrast, Finngen, Biobank Japan and iPSYCH required more passive participant involvement with no or little study engagement, as samples were collected from routine biospecimens or during clinical visits. We observed significant heritability of sex only in the studies that required more active participation (h^2^ on liability scale=3.0% (P=3×10^−127^) and 2.3% (P=2×10^−14^), in 23andMe and UK Biobank, respectively), while no significant heritability was detected in the three more passive studies (**Figure 2** and **Supplementary Table 4**).

**Figure 2:**
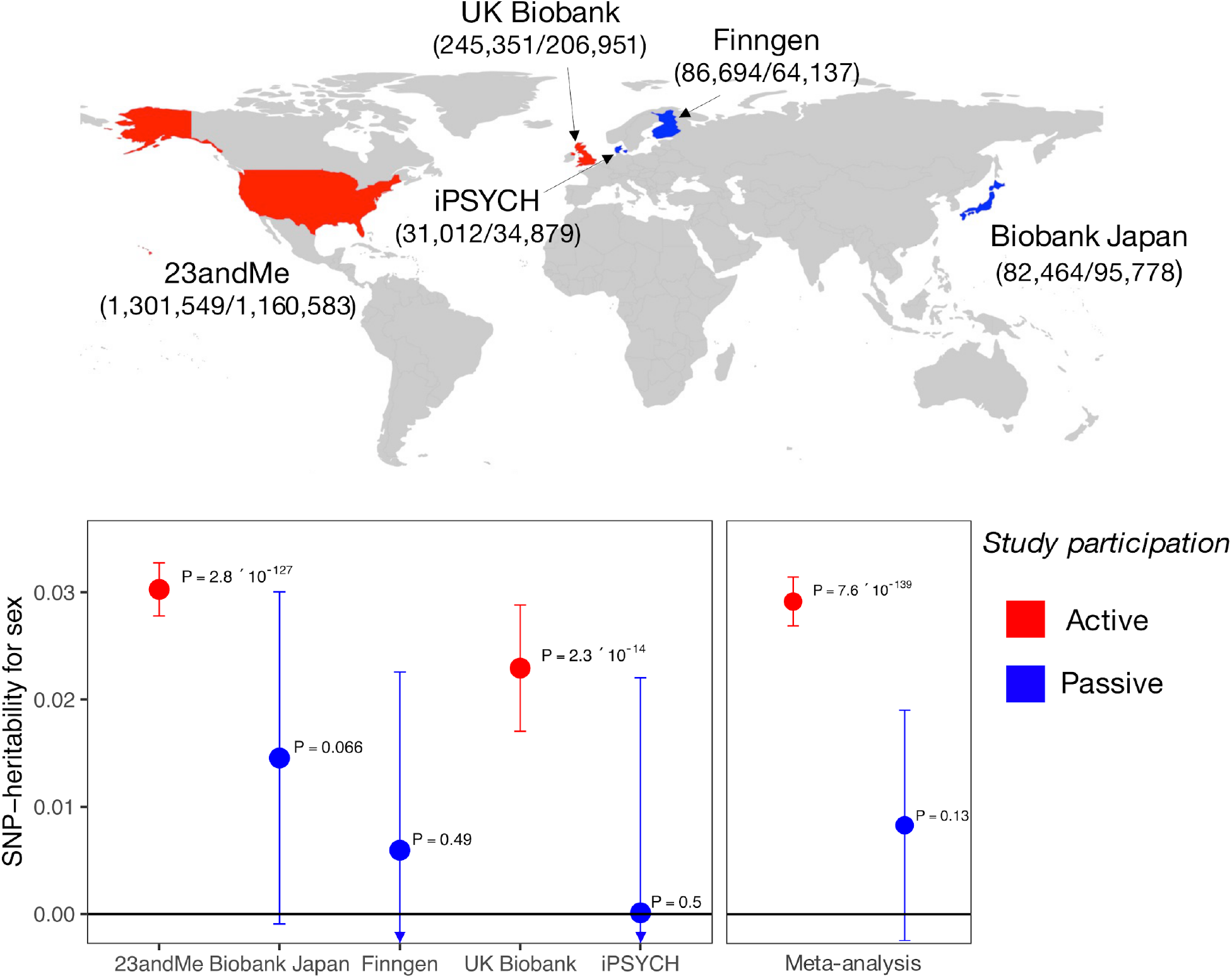
SNP-heritability on the liability scale for sex across 5 studies (23andMe: N=2,462,132; Biobank Japan: N=178,242; Finngen: N=150,831; UK Biobank: N=452,302; iPSYCH: N=65,891). Error bars represent the confidence interval for the SNP-heritability estimate. For each study, we report in parentheses the number of females/males included in the analysis. In red studies characterized by “active” participation, in blue studies with “passive” participation. iPSYCH heritability is negative and therefore set to 0. Definitions of “active” and “passive” are ad-hoc for this study and encompass heterogeneous enrollment strategies and consent modalities.

iPSYCH, in particular, showed the lowest heritability estimate, consistent with its study design that retrieved routinely collected neonatal dried blood spots from a random sample of individuals born between 1981 and 2005, who were alive and resident in Denmark on their first birthday, thus minimizing both participation and survival bias. In aggregate, these findings suggest that many autosomal signals for sex represent underlying mechanisms that influence sex-differential study participation rather than sex-differential pre-sampling mortality. We do not preclude the possibility that a small number of loci might influence sex-differential survival from *in utero* to age 30 years, which should be explored in future studies of younger individuals.

To demonstrate the statistical basis of our observed sex-differential participation bias, we simulated a phenotype that is uncorrelated with sex and has a heritability of 30% in 350,000 individuals, half males and half females (**Figure 3A)**. Under different sampling scenarios, we found that sex becomes significantly heritable on autosomes if study participation is dependent on the phenotype in a sex-differential manner (**Figure 3B**). In the presence of this bias, autosomal variants associated with the phenotype are also associated with sex in a dose-response manner. As a consequence, Mendelian randomization (MR) analysis would wrongly identify a causal relationship between the phenotype and sex (**Figure 3C)**. An alternative explanation for our findings is that sex is a causal factor for the phenotype that influences study participation (**Extended Figure 1 panel A**) or that both sex and the phenotype drive participation independently (**Extended Figure 1 panel B**); however, we show using both real data and simulations that these models are highly unlikely (see **Supplementary Note**).

**Figure 3:**
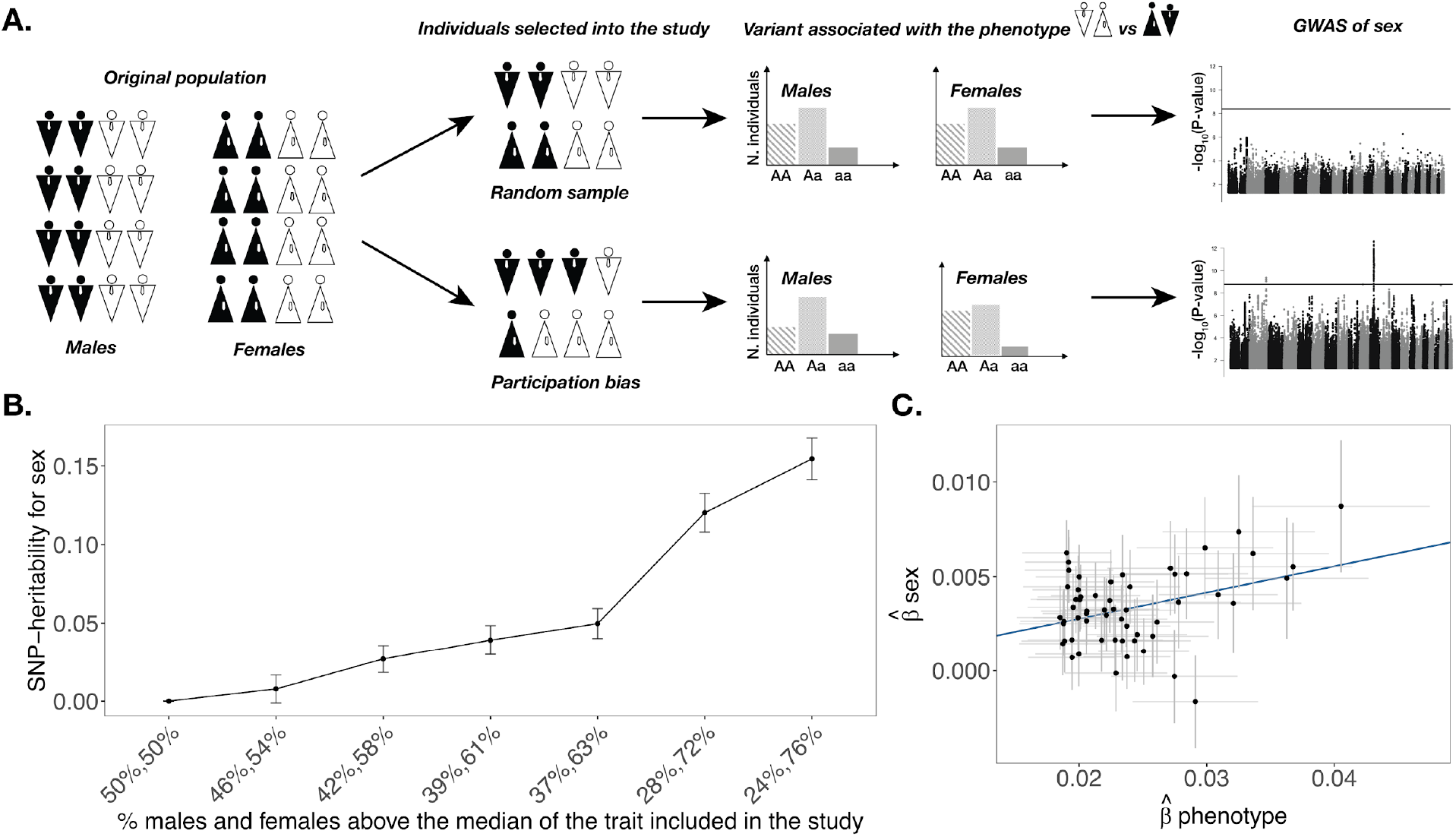
Illustration of the concept and consequences of sex-differential participation bias. **A.** Schematic representation of sex-differential participation bias. Because males and females distribute differently for a certain trait in the selected study population, variants associated with the trait become associated with sex. **B.** heritability of sex increases as function of sex-differential participation bias expressed as the percentage of males and females above the median of the phenotype included in the study (N=350,000). If there is no bias this value is 50% for both males and females. Dots represent the SNP effect size, error bars represent the confidence intervals for the heritability estimate**. C.** variants associated with the phenotype are also associated with sex in a dose-dependent manner. Mendelian randomization would indicate a causal relationship between sex and the phenotype. Here we consider only variants genome-wide significantly associated with the phenotype in the fourth scenario of panel B (39%,61%). Dots represent the SNP effect size, error bars represent the confidence intervals for the SNP effect size.

### Genetic analyses reveal determinants of sex-differential participation bias

We then systematically tested complex traits for evidence of a shared genetic architecture with sex-differential participation bias in UK Biobank and 23andMe. By analysing summary data from 4,155 publicly available GWASs^20^, we showed that sex-associated signals are enriched for pleiotropic associations (P<2×10^−16^; Chi-square test comparing sex-associated SNPs *vs* all SNPs); half of the genome-wide significant imputed signals for sex were associated with at least one complex trait, and one-fifth were associated with five or more traits (**Supplementary Table 5**). Genetically correlated traits spanned a diverse range of health outcomes, including blood pressure, type 2 diabetes, anthropometry, bone mineral density, auto-immune disease, personality traits and psychiatric diseases.

Genome-wide autosomal correlation analyses with 38 health and behavioral traits highlighted 22 significant associations with sex in 23andMe and 5 in UK Biobank (**Figure 4** and **Supplementary Table 6**). We noted that the genetic correlates of sex overlapped only partially between 23andMe and UK Biobank (*rg*=0.50, P-value=4×10^−34^), which was reflected in several trait-specific study discordant associations. For example, higher educational attainment was associated with female sex in UK Biobank (rg=0.25, P=7×10^−12^), while the opposite direction of association was observed in 23andMe (*rg*=−0.31, P=9×10^−81^). This finding demonstrates that the determinants of sex-differential participation bias may vary substantially between studies.

**Figure 4:**
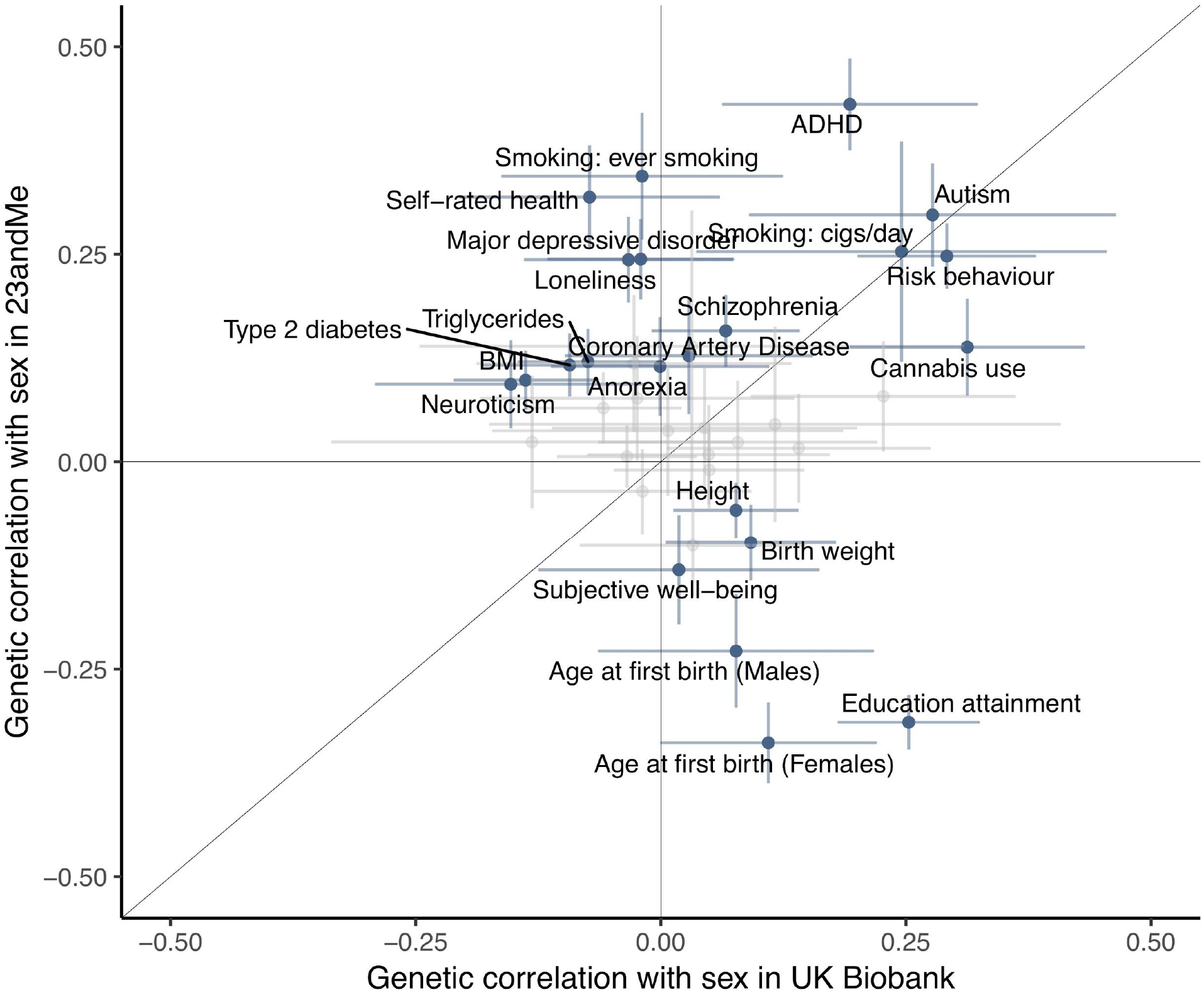
Genetic correlation with being born female *vs* male and 38 traits in UK biobank and 23andMe. Only correlations that are significant in at least one of the two studies are highlighted. Dots represent the genetic correlation estimates, error bars represent the confidence intervals.

A notable autosomal signal for sex was at the obesity-associated *FTO* gene locus, where the body mass index (BMI) increasing allele was observed in 23andMe at higher frequency in males compared to females (rs10468280, OR 1.02 [1.02-1.03], P=4.4×10^−36^ **Supplementary Table 1**). The same direction and magnitude of effect at the *FTO* locus was also observed in UK Biobank (OR= 1.02 [1.01-1.03], P=3.6×10^−5^), and subsequent Mendelian randomization analyses supported a causal effect of BMI on sex in both 23andMe and UK Biobank (**Supplementary Table 7**). We note however that there was considerable heterogeneity in the dose-response relationship between BMI variants and sex, and it remains unclear how genetically higher BMI leads to sex-differential study participation. Intriguingly the genetic correlation between BMI and sex was discordant between UK Biobank (rg=−0.13, P=2×10^−04^) and 23andMe (rg=0.10, P=9×10^−08^), and this difference between studies appeared attributable to negative confounding by educational attainment (**Supplementary Table 7**). These results reinforce the need for caution when inferring causality from genetic correlations.

Traditional approaches to identify study participation bias compare the distribution of a phenotype in the study with that of a representative population. Using this approach, we confirmed our genetic inference that the difference in education level between UK Biobank participants and UK census data was larger in females than in males (**Figure 5A** and **Supplementary Table 8**). Such greater differential participation by education among females can also be observed, without the need for census data, by comparing the distribution of polygenic scores for education between males *vs* females. If we had a completely representative sample, we would not expect any differences in the distribution of the polygenic score for educational attainment between males and females (i.e. all the differences in measured educational attainment between the two sexes are expected to be due to environmental factors). Any difference in the polygenic score distribution needs therefore to be explained by selection acting on educational attainment which is either determined by sex or it has occurred differentially between men and women.

**Figure 5:**
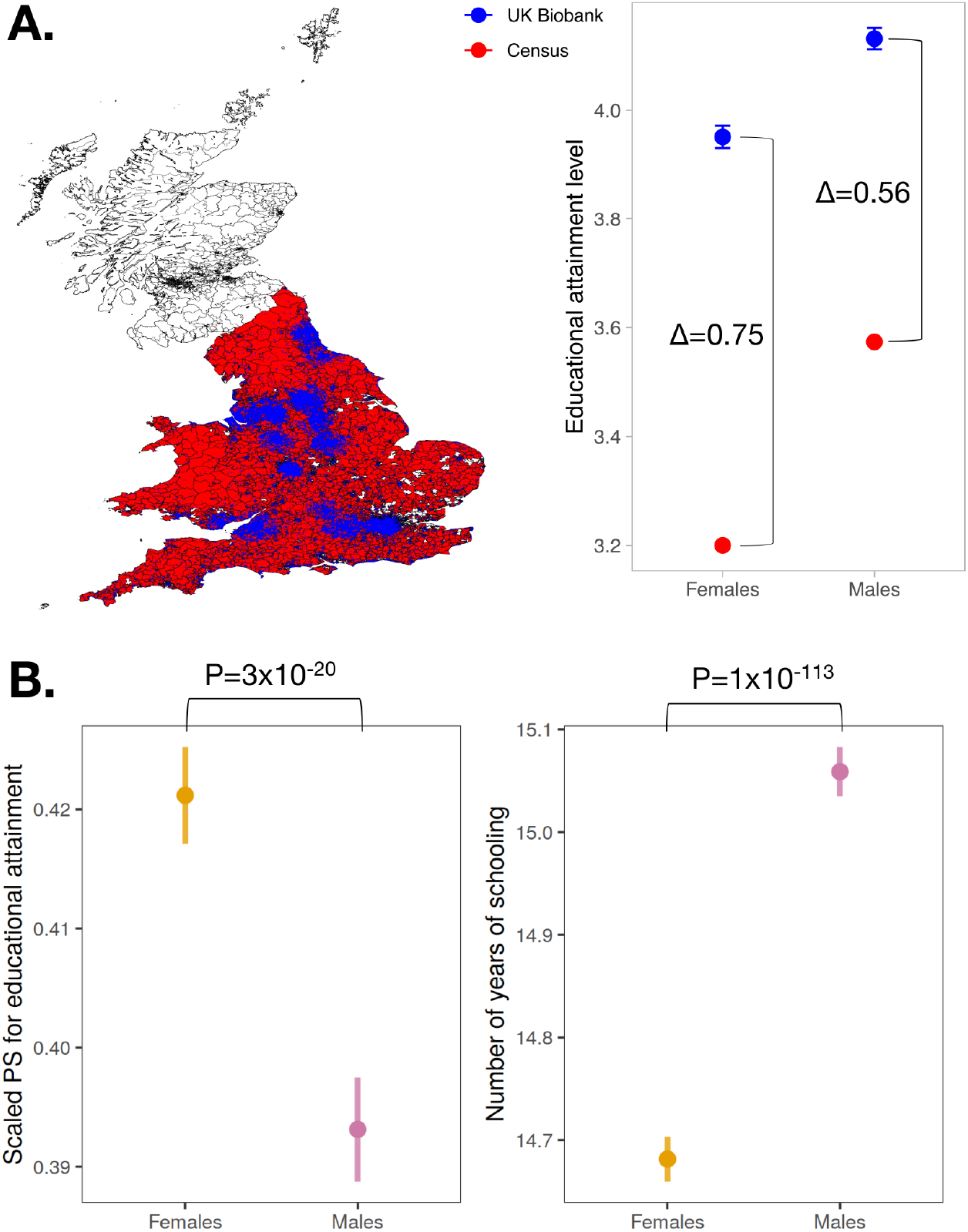
**A.** Comparing highest education level between 2011 England and Wales census data (in red, N=29,492,209) with UK Biobank (in blue, N=411,845). We only considered regional census districts with at least one UK Biobank participant. The difference in the average education level between males and females is higher in the general population than in participants in UK Biobank. Dots represent the mean taking into account the sampling design, error bars represent the confidence intervals. No confidence intervals were considered for the census data because the entire population was included. **B.** Polygenic score (PS) for educational attainment is significantly higher in females (N=194,282) compared to males (N=167,219) in UK Biobank, vice versa, the number of years of schooling is higher in males (two-sided t-test). Dots represent the mean value, error bars represent the standard deviation.

By using data from the SSGAC consortium^21^, which did not include UK Biobank or 23andMe, we constructed a polygenic score for educational attainment. We first examined iPSYCH, where we don’t expect participation bias and indeed we saw no significant differences in the distribution of the polygenic score for educational attainment between males and females (P=0.78). In UK Biobank, the mean polygenic score was higher in females than in males (P=7×10^−23^; t-test), consistent with the census data comparison. We note that, opposite to the polygenic score, the reported education level in UK Biobank is significantly higher in males compared to females (t-test P=1×10^−113^) (**Figure 5B**). Therefore, on its own, the distribution of the phenotype among study participants does not inform the direction and degree of sex-differential participation bias.

Educational attainment is one of few traits for which representative data are available, via the UK census. For other traits, where such information is not collected, genetic analysis in the form of polygenic scores provides a unique opportunity to identify novel sex-differential determinants of study participation.

### Sex-differential participant bias can influence downstream genetic analyses

Next, we illustrated the potential effects of sex-differential participation bias on downstream genetic analyses using simulated and empirical data (**Extended Figures 3–4, Supplementary Figures 2–5, Supplementary Note and Supplementary Table 9-10**).

First, we performed simulation analyses to demonstrate that this bias can lead to spurious genetic correlations between two traits by exacerbating or attenuating the effects of overall participation bias (**Extended Figure 3**). Furthermore, it can lead to an incorrect causal inference (in MR analyses) between two phenotypes in a sex-differential manner (**Extended Figure 4)**. For example, Censin and colleagues recently described sex differences in the causal effect of BMI on cardiometabolic outcomes in UK Biobank^22^. They concluded that the magnitude of increase in risk for Type 2 diabetes (T2D) due to obesity differs between males and females. We attempted to confirm their results in light of our observations, and found that their findings were likely biased due to reasons other than sex-differential participation bias (see **Supplementary Note**). However, we demonstrate through simulation analyses that sex-differential participation bias could indeed lead to incorrect inferences in such Mendelian randomization analyses (**Supplementary Table 10**). With only modest BMI-related sex-differential participation bias, we saw artificial sex differences in the association between a BMI genetic score and T2D and, in the most extreme sampling parameters, the direction of sex difference was swapped, with BMI genetic score-T2D effect estimates ranging from OR_male_=2.71 and OR_female_=3.49 to OR_male_=3.86 and OR_female_=2.61. These results highlight the challenges of performing and interpreting sex-specific analyses in studies where the exposure variable may be influenced by sex differences in participation bias.

Second, in a scenario where sex-differential participation exists, adjusting for sex as a covariate in a GWAS could bias effect estimates of any genetic analysis (**Supplementary Figure 3)**. To explore this possibility, we performed 565 GWASs of heritable traits in the UK Biobank and estimated the genetic correlation between each trait with and without inclusion of sex as a covariate. The results were highly consistent (**Supplementary Figure 4**) between the two models, with sizable differences (indicated by lower genetic correlations) observed only for highly sex-differentiated traits (e.g. testosterone levels). Importantly, sex-differential participation bias did not impact the genetic correlation between males and females for each phenotype (**Supplementary Figure 5)**. We caution that, although inclusion of sex as covariate did not seem to impact most traits in these analyses, this issue might lead to significant differences between models as sample sizes continue to grow.

## Discussion

Most large-scale biobank studies are not designed to achieve cohorts who are accurately representative of the general population^23,24,25,26,27,28^. Lack of representation is not *per-se* problematic if this is considered when interpreting study findings^6^. Here, we show an example of how sex-differential study participation bias could lead to spurious associations and ultimately incorrect biological inferences. In practice, the impact of differential participation bias on genetic results is hard to tease apart for most traits. We used sex, which provides a robust negative control as it has no autosomal determinants, to identify determinants of study participation bias that differentially impact males and females.

We demonstrate that sex-differential participation bias results in sex showing spurious heritability on the autosomes and being genetically correlated with the complex traits that underlie such bias. This is of course highly important in iPSYCH and other studies of psychiatric disorders and traits strongly associated with gender as, e.g., autism, ADHD, and depression, but the implications generalize to many other risk factors and phenotypes. For example, alleles associated with higher BMI are under-represented in females compared to males in both UK Biobank and 23andMe. This suggests that females with higher genetic susceptibility to obesity are less likely to participate in studies than their male equivalents (or that genetically lean males are more likely to), although the mechanism by which genetically determined BMI influences non-participation is unclear. These sex-differential biases may also have directionally opposite effects between studies - alleles associated with higher educational attainment were under-represented in 23andMe females but over-represented in UK Biobank females. While these results reflect differences in participation between men and women, we do not yet understand the *mechanisms* by which differences in BMI or education lead to differential participation between sexes. This may be due to clinical, social or cultural factors that lead to changes in the perception or expectations of individuals when deciding to engage in research studies. Our results are consistent with the larger effect - and larger bias - observed for the association between sex and cardiovascular mortality when UK Biobank is compared to a representative health survey^29^. We conclude that sex-differential participation can induce false sex-differential associations (or obscure true associations) and complicate the study of health disparities between males and females.

While study design and participant recruitment strategy are the most likely factors influencing participation bias, we showed that both novel and existing methods can be applied to reduce the impact of such bias. Inverse-probability-of-sampling-weighted (IPW) regression has been applied to achieve unbiased estimates from analyses of case-control data^30,31^. Dudbridge, Mahmoud and colleagues^32,33^ proposed a correction for selection that occurs when performing case-only analyses. However, the same technique can correct for selection that is conditioned on any trait as long as GWAS can be performed on it. We propose two additional conceptual frameworks and show how they can be implemented in genomicSEM^34^. First, we developed an application of Heckman correction for genetic data. Heckman correction^35^ is commonly used in econometrics to correct for the association between an exposure X and outcome Y when the outcome is observed only in study participants and thus is subject to participation bias. The intuition behind Heckman correction is that the predicted probability of study participation (S) can be used to adjust the association between Y and X.

Second, we propose a novel method that is based on the following intuition: the magnitude of participation bias introduced between X and Y under selection is proportional to their effects on the probability of study participation (S). By specifying a model where the bias and the effect that introduces the bias are forced through a single path, the correct genetic correlation between Y and X can be retrieved from the GWAS of Y and X in the selected samples and S. This method, unlike Heckman correction, does not require the predicted probability of study participation, but instead a GWAS of participating individuals versus the population is sufficient. Details of both of these methods are provided in the **Supplementary Note**.

While we validated the two approaches via simulations (**Supplementary Table 11**), future work is needed to apply these methods to examples in real data. The biggest challenge to the implementation of both approaches to bias correction is that they require unbiased estimates of allele frequencies in the target population. The generation of such information, for example by establishing a “census of human genetic variation”, should be the primary focus of future activities in this area. Some extremely large genomic databases exist, such as gnomAD^36^. However, these are unlikely to be representative due to inclusion of data from studies with a wide range of designs and settings. Where legislation allows, designs such as used by iPSYCH could be implemented^5^. That study has already shown the value of generating accurate population-based estimates of rare copy number variants^37^. A future study much smaller than iPSYCH could valuably inform population allele frequencies using neonatal dried blood spots in a manner that protects anonymity, while significantly strengthening the inferences derived from other larger non-representative studies. Such an approach would be necessary to implement the bias correction frameworks proposed above.

In summary, we demonstrate that genetic analyses can uniquely profile the complex traits and behaviours that contribute to participation bias in epidemiological studies. We hope that future studies will build on these findings to create resources and tools that more systematically identify and correct for broader forms of participation bias and their effects on genetic association results.

## Acknowledgments

We acknowledge Prof. George Davey Smith for insightful comments. This research was conducted by using the UK Biobank Resource under application 31063. AG was supported by the Academy of Finland Fellowship (323116). This work was supported by the Medical Research Council (Unit Programme number MC_UU_12015/2).

MGN is a fellow of the Jacobs Foundation, is supported by ZonMW grants 849200011 and 531003014 from The Netherlands Organisation for Health Research and Development & a VENI grant awarded by NWO (VI.Veni.191G.030).

AG has received support by Academy of Finland Fellow grant N. 323116.

The FinnGen project is funded by two grants from Business Finland (HUS 4685/31/2016 and UH 4386/31/2016) and eleven industry partners (AbbVie Inc, AstraZeneca UK Ltd, Biogen MA Inc, Celgene Corporation, Celgene International II Sàrl, Genentech Inc, Merck Sharp & Dohme Corp, Pfizer Inc., GlaxoSmithKline, Sanofi, Maze Therapeutics Inc., Janssen Biotech Inc). Following biobanks are acknowledged for collecting the FinnGen project samples: Auria Biobank (www.auria.fi/biopankki), THL Biobank (www.thl.fi/biobank), Helsinki Biobank (www.helsinginbiopankki.fi), Biobank Borealis of Northern Finland (https://www.ppshp.fi/Tutkimus-ja-opetus/Biopankki/Pages/Biobank-Borealis-briefly-in-English.aspx), Finnish Clinical Biobank Tampere (www.tays.fi/en-US/Research_and_development/Finnish_Clinical_Biobank_Tampere), Biobank of Eastern Finland (www.ita-suomenbiopankki.fi/en), Central Finland Biobank (www.ksshp.fi/fi-FI/Potilaalle/Biopankki), Finnish Red Cross Blood Service Biobank (www.veripalvelu.fi/verenluovutus/biopankkitoiminta) and Terveystalo Biobank (www.terveystalo.com/fi/Yritystietoa/Terveystalo-Biopankki/Biopankki/). All Finnish Biobanks are members of BBMRI.fi infrastructure (www.bbmri.fi).

## Author contributions

**Study design:** NP, MC, PN, CC, MDZ, AbA, DH, BMN, RKW, MGN, JRBP, AG. **Data Analysis:** NP, MC, PN, GM, AbA, BH, MK, VMR, PDBP, MB, JK, TDA, MDZ, RB, ADB, AA, DH, MGN, JRBP, AG. **Results interpretation:** NP, MC, AbA, CC, FRD, KKO, RB, PJ, BMN, RKW, MGN, JRBP, AG. **Provided Data:** PN, AbA, VMR, TDA, TM, EG, YO, ADB, AA, DH, MBN, MGN, JRBP, AG. **Wrote the Manuscript:** NP, MC, BMN, MGN, JRBP, AG.

## Materials and Methods

### Contributing cohorts GWAS

Genome wide association was conducted in 5 different cohorts: 23andMe, Uk biobank, iPSYCH, FinnGen and Biobank Japan, for a total of 3,309,398 samples: 1747070 female and 1562328 male.

Full detailed cohort description, recruitment and genotyping information can be found in the supplementary materials. For all GWAS analyses females were coded as 1 while male as 0.

#### 23andMe

##### Ancestry assignment

We restrict participants to a set of individuals who have a specified ancestry determined through an analysis of local ancestry. Briefly, our algorithm first partitions phased genomic data into short windows of about 300 SNPs. Within each window, we use a support vector machine (SVM) to classify individual haplotypes into one of 31 reference populations (https://www.23andme.com/ancestry-composition-guide/).

The SVM classifications are then fed into a hidden Markov model (HMM) that accounts for switch errors and incorrect assignments, and gives probabilities for each reference population in each window. Finally, we used simulated admixed individuals to recalibrate the HMM probabilities so that the reported assignments are consistent with the simulated admixture proportions. The reference population data is derived from public datasets (the Human Genome Diversity Project, HapMap, and 1000 Genomes), as well as 23andMe customers who have reported having four grandparents from the same country. European participants were identified using the following classification probabilities: European + Middle Eastern > 0.97, European > 0.90.

##### Relatedness

A maximal set of unrelated individuals was chosen for each analysis using a segmental identity-by-descent (IBD) estimation algorithm^38^. Individuals were defined as related if they shared more than 700 cM IBD, including regions where the two individuals share either one or both genomic segments IBD. This level of relatedness (roughly 20% of the genome) corresponds approximately to the minimal expected sharing between first cousins in an outbred population.

##### Association

We compute association test results for the genotyped and the imputed SNPs. For case-control phenotypes, we compute association by logistic regression assuming additive allelic effects. For tests using imputed data, we use the imputed dosages rather than best-guess genotypes. As standard, we include covariates for age, the top five principal components to account for residual population structure, and indicators for genotype platforms to account for genotype batch effects. The association test P-value we report is computed using a likelihood ratio test, which in our experience is better behaved than a Wald test on the regression coefficient.

For QC of genotyped GWAS results, we excluded SNPs that were only genotyped on our “v1” and/or “v2” platforms due to the small sample size. Using trio data, we excluded SNPs that failed a test for parent-offspring transmission; specifically, we regressed the child’s allele count against the mean parental allele count and flagged SNPs with fitted β<0.6 and P<10^−20^ for a test of β<1. We excluded SNPs with a Hardy-Weinberg P<10^−20^, or a call rate of <90%. We also tested genotyped SNPs for genotype date effects and flagged SNPs with P<10^−50^ by ANOVA of SNP genotypes against a factor dividing genotyping date into 20 roughly equal-sized buckets. We excluded SNPs with a large sex effect (ANOVA of SNP genotypes, r^2^>0.1). Finally, we excluded SNPs with probes matching multiple genomic positions in the reference genome (‘self chain’). For imputed GWAS results, we excluded SNPs that had strong evidence of a platform batch effect. The batch effect test is an F test from an ANOVA of the SNP dosages against a factor representing v4 or v5 platform; we flagged results with P<10^−50^. Variants with imputation INFO score < 0.8 or MAF < 0.01 were excluded from the analysis.

#### UK Biobank

##### Ancestry assignment

We defined a subset of ‘white European’ ancestry samples using a k-means-clustering approach that was applied to the first four principal components calculated from genome-wide SNP genotypes. Individuals clustered into this group who self-identified by questionnaire as being of an ancestry other than white European were excluded.

##### Association

Association testing was performed using a linear mixed model implemented in BOLT-LMM^39^ v2.3.4 to account for cryptic population structure and relatedness. Only autosomal genetic variants that were common (minor allele frequency (MAF) > 1%), passed quality control in all 106 batches and were present on both genotyping arrays were included in the genetic relationship matrix. Genotyping chip, age at baseline and ten genetically derived principal components were included as covariates. Variants with imputation INFO score < 0.8 or MAF < 0.01 were excluded from the analysis.

#### iPSYCH

##### Relatedness

In total 78,050 genotyped individuals were available for analysis. Among them, 11,128 individuals were identified to be related (piHAT > 0.2). Relatedness was measured using Identity by descent (IBD) analysis using Plink V. 1.90. Those individuals with piHAT > 0.2 were considered as related. Among the 11,128 related individuals, one of each pair of related individuals was removed randomly (retaining 5,476 individuals). Hence, a total of 72,398 individuals were taken forward for principal component analysis.

##### Ancestry assignment

Principal component analysis was performed across 72,398 individuals using R package SNPRelate v1.20^40^. Only high quality imputed variants (N markers=22,576) with MAF > 0.01, missing rate < 0.01 and INFO > 0.8 were included in the analysis. The variants were LD pruned (R2 < 0.2) before PCA. All the pairwise scatter plots for PCs 1 to 10 were visualized and the first 3 PCs were chosen for outlier detection. Among the 72,398 individuals, 44,158 individuals were Danes for at least three generations (they, their parents, their paternal and maternal grandparents, all born in Denmark). A 3-dimensional Ellipsoid was constructed using the principal components 1, 2 and 3 of only the pure Danes with a radius of 5 standard deviations. In total 6,499 individuals were outside this ellipsoid and subsequently excluded as so population outlier (leaving 65,899 individuals).

##### Association

Among the 65,899 individuals, information about sex was missing for eight individuals (either NA or mismatched during cross-verification), hence they were removed leaving behind 65,891 for GWAS analysis, which comprises 31,012 females and 34,879 males. The 65,891 individuals included cases of six psychiatric disorders and 22,439 controls (without any of the six psychiatric disorders).

The GWAS analyses were conducted using Plink V1.90 using --dosage argument. The covariates included were: age, age squared, 10 first principal components, wave number (as one-hot encoding), psychiatric disorder type (as one-hot encoding). After the GWAS, the variants with INFO < 0.8 and MAF < 0.01 are removed. Analysis performed only in controls gave similar results, but with larger standard errors due to the smaller sample size.

#### Finngen

##### Ancestry assignment

For principal components analysis, FinnGen data was combined with 1000 genomes data. Related individuals (<3rd degree) were removed using King software^41^. We considered common (MAF >= 0.05) high quality variants: not in chromosome X, imputation INFO>0.95, genotype imputed posterior probability>0.95 and missingess<0.01. LD-pruned (r^2^<0.1) common variants were used for computing PCA with Plink 1.92.

##### Association

SAIGE mixed model logistic regression (https://github.com/weizhouUMICH/SAIGE/releases/tag/0.35.8.8) was used for association analysis. Age and 10 PCs and genotyping batch were used as covariates. Each genotyping batch was included as a covariate for an endpoint if there were at least 10 cases and 10 controls in that batch to avoid convergence issues. Variants with imputation INFO score < 0.8 or MAF < 0.01 were excluded from the analysis.

##### Biobank Japan

We retrieved individuals’ sex from medical records, and excluded samples who have inconsistent sex with genetically determined sex. In total, we used 95,778 males and 82,464 females for analysis. GWAS was conducted using PLINK v2.0 under a linear regression model with covariates including age and top 20 PCs. Variants with imputation INFO score < 0.8 or MAF < 0.01 were excluded from the analysis.

### Identification of independent loci and additional QC of results from 23andMe

To evaluate if our sex-associated genome-wide significant signals were attributable to technical artifacts we embarked in additional quality controls. First, we used the FUMA v1.3.5d pipeline^42^ to identify independent loci. In particular, we used pre-calculated LD (linkage disequilibrium) structure based on the European 1000 Genome panel to identify genome-wide significant SNPs independent from each other at r^2^<0.6.

If LD blocks of independent significant SNPs are closely located to each other (< 250 kb based on the most right and left SNPs from each LD block), they are merged into one genomic locus. FUMA also identifies independent *lead* SNPs within a locus if they are independent of each other at r^2^ <0.1. Each genomic locus can thus contain multiple independent significant SNPs and lead SNPs. This approach resulted in 158 loci which are reported in **Supplementary Table 1**. For each locus, we identified one directly genotyped SNP with P-value < 5×10^−8^. This resulted in 78 SNPs since not all loci had a genome-wide significant directly genotyped SNP. We extracted 50 *bp* upstream and downstream of each SNP using h19 reference genome and the R function *getSeq* from the package BSgenome 1.58.0. We chose 50 *bp* as this is the probe length on the Illumina Global Screening array. We used BLAT v.407 (https://genome.ucsc.edu/cgi-bin/hgBlat?command=start) to search each extracted sequence vs the human genome. We considered only matches on chromosome X and Y with 95% or greater similarity. We also considered stricter quality control metrics: Hardy-Weinberg disequilibrium threshold > 1×10^−6^, MAF > 5% and call rate > 98%.

All downstream analyses looking at the aggregate effect of variants across the genome are done using all the variants that passed cohort-specific quality controls without considering the strict quality controls thresholds described above.

### Pleiotropy analysis

To test the relevance of our sex associated signals with other traits we used the results from the analysis of Watanabe *et al*^20^, which considered GWAS results from 4,155 publicly available GWASs. For each locus we count the number of associated traits and categorized as 0, 1, 2, 3, 4, 5+. These results can be obtained by combining results from Supplementary Table 4 of Watanabe *et al* together with all the SNPs tested for pleiotropy, which are available here: https://github.com/dsgelab/genobias. We then use a chi-square test to compare the count distribution for the number SNPs that were GW-significant associated with sex *vs* all SNPs considered by Watanabe *et al*.

### Extract results from the GWAS catalog

We considered the most significant SNP for each of the 158 genome-wide significant loci and extracted all the SNPs in LD (r^2^ > 0.2 and distance < ±500Mb). To extract these SNPs we used the R implementation of LDproxy (https://ldlink.nci.nih.gov/?tab=ldproxy) and used an LD reference panel from 1000 genomes Northern Europeans. To identify traits significantly associated with these proxy SNPs we interrogate the GWAS catalog^43^ using the R package gwascat v. 2.22.0. The GWAS catalog was extracted in date 2^nd^ December, 2019. We only considered reported association with P < 5×10^−8^ and extracted the EFO terms.

### Comparison of full GWAS vs individuals < 30 years old in 23andMe

To identify loci significantly associated with sex in individuals younger than 30 years old at recruitment we used the same pipeline described in “Identification of independent loci and additional QC of results from 23andMe”. To test the difference in effect sizes between the two analysis we used the following test:

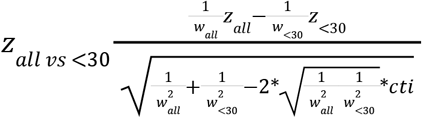

Where 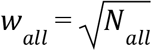 where N_all_ is the full sample size and 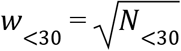 where N_<30_ is the sample size for the people younger than 30. *z*-scores *z_all_* and *z*_<30_ are obtained from the corresponding GWAS results, and *cti* is the intercept from the LD-score genetic correlation between the two analyses. We can obtain *z*-scores for the difference between the two analyses reweighted by the corresponding sample size to allow for differences in sample sizes between the two analyses. The test is analogous to the test for a sum of z statistics forms dependent GWAS as presented in Baselmans et al.^44^ and Jansen et al^45^, and similar to the method used by Nolte et al^46^.

In order to verify if sample overlap would affect our results, we derived the expected *z*-scores for the GWAS run without the samples with age < 30. This was estimated as:

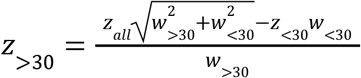

Where z_>30_ is the expected *z*-score in people older than 30, and 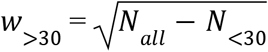. Differences tested between the >30 and <30 datasets showed no difference with the ones observed in the overall dataset.

### Heritability estimation of sex

We used LD-score regression^47^ to estimate the proportion of variance in liability to sex at birth that could be explained by the aggregated effect of the SNPs. The method is based on the idea that an estimated SNP effect includes the effects of all SNPs in LD with that SNP. On average, a SNP that tags many other SNPs will have a higher probability of tagging a causal variant than a SNP that tags few other SNPs. Accordingly, for polygenic traits, SNPs with a higher LD-score have on average stronger effect sizes than SNPs with lower LD-scores. When regressing the effect size obtained from the GWAS against the LD-score for each SNP, the slope of the regression line gives an estimate of the proportion of variance accounted for by all analyzed SNPs. We included 1,217,312 SNPs (those available in the HapMap 3 reference panel). We used stratified LDscore regression, including LD and frequency annotation, similar to what is used by Gazal et al.^48^ since this has been shown to reduce bias in heritability estimation^49,50^.

Since sex is a dichotomous trait, which frequency changes across studies, we have transformed the observed heritability 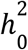 into liability scale 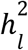 using the following formula^51^:

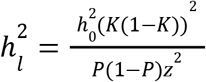

Where *K* is the prevalence of sex in the population (50%), *P* is the proportion of females in the study and *z* is the height of the normal curve corresponding to the prevalence of sex in the population.

For estimation of heritability in Japan Biobank we used a LD score reference panel based on East Asian participants in 1000 genomes.

### Genetic correlations

We used cross-trait LD-score regression to estimate the genetic covariation between traits using GWAS summary statistics^28^. The genetic covariance is estimated using the slope from the regression of the product of z-scores from two GWAS studies on the LD-score. The estimate obtained from this method represents the genetic correlation between the two traits based on all polygenic effects captured by SNPs. Standard LD-scores were used as provided by Bulik-Sullivan et al^52^ based on the 1000 genomes reference set, restricted to European populations.

The decision of which summary statistics to include in the genetic correlation analysis was taken before analyzing the data by consensus across the authors of the paper.

### MR analysis and genomicSEM regression for BMI and sex

We tested for possible causal effects of BMI on sex, induced by sex-differential participation bias, in both 23andMe and UK Biobank through MR. As instruments for the exposure, we utilized the 97 index SNPs associated with BMI reported by Locke and colleagues^53^. We tested different methods (MR Egger, Weighted median, Inverse variance weighted, Simple mode, Weighted mode) as implemented in the R package TwoSampleMR^54^.

We then further investigate whether the discordance in genetic correlations between BMI and sex in UK Biobank (rg=-0.13, P=2×10^−04^) and 23andMe (rg=0.10, P=9×10^−08^) is due to a confounding effect of educational attainment. By using the respective GWAS summary statistics, we fitted the following multiple regression model in genomicSEM^34^ to estimate the genetic correlation between BMI and sex controlling for educational attainment:

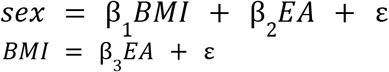

Results for both analyses are reported in **Supplementary Table 7**.

### Generation of genetic scores for educational attainment

We used summary statistics for a GWAS of years of education^21^, which did not include Uk Biobank and 23andMe, to construct the polygenic score. This score was generated using *PRSice v.2.0*^55^. Briefly, PRSice performs a pruning (distance=250KB and r^2^=0.1) and thresholding approach. We then selected the P-value threshold that maximizes the r2 between the score and educational attainment in UK Biobank (P=0.195, N.SNPs=39,014). The polygenic score was only constructed for a subset of the UK Biobank containing white-British unrelated individuals (N=361,501) as described here: https://github.com/Nealelab/UK_Biobank_GWAS

We constructed the polygenic score on the dataset including both males and females and then we compared if the average polygenic score differed between males and females using a *t-test*. Next, we compared the average years of education in the same dataset. We recorded the education level variable in Uk Biobank (“6138”) into years of education following the approach used by the SSGAC consortium: 1=20 years; 2=15 years; 3=13 years; 4=12 years; 5=19 years; 6=17 years; −7=6 years; −3=missing.We then test for significant differences in education between males and females using a *t-test*.

### Census data analysis

We obtained information about educational attainment from the UK Census from the year 2011. Data were extracted from the Office for National Statistics: https://www.nomisweb.co.uk/census/2011. We coded the qualification level collected in the census to match the corresponding levels in UK Biobank:

#### Census

No qualifications => 1
Level 1 qualifications => 2
Level 2 qualifications => 3
Apprenticeship => 4
Level 3 qualifications => 5
Level 4 qualifications and above: 6
Other qualifications => NA

#### UK Biobank

1: College or University degree => 6
2: A levels/AS levels or equivalent => 5
3: O levels/GCSEs or equivalent => 2.5
4: CSEs or equivalent => 2.5
5: NVQ or HND or HNC or equivalent => 6
6: Other professional qualifications eg: nursing, teaching => 6
-7: None of the above => 1
-3: Prefer not to answer => NA

Information from the 2011 census was grouped by three age bins (35-49, 50-64, 65+), sex and Middle Layer Super Output Area (MSOA) regions from England and Wales. In total, 6,050 MSOA regions with at least one UK Biobank participant were included. To map each individual to an MSOA region we used the home location coordinates (variables 22702 and 22704) with the moving date that was closest to 2011. We then use the *sp* v. 1.4-5 R package (*over* function) to map the coordinates to the MSOA region coordinates obtained from https://census.mimas.ac.uk/dataset/2011-census-geography-boundaries-middle-layer-super-output-areas-and-intermediate-zones-7. To estimate the average education level, separately in men and women in UK Biobank and in the census, we use the *svydesign* function from the survey *v*. 4.0 R package. This function implements different types of sampling designs and, in this analysis, we used a stratified sampling design with three strata: age, sex, and MSOA region.

## Data availability

GWAS results are available through GWAS catalogue accession numbers GCST90013473 (23andMe) and GCST90013474. Full summary statistics for 23andMe are available upon request from https://research.23andme.com/dataset-access/.

**Code Availability**

Scripts are available at: https://github.com/dsgelab/genobias

Collaborators

## iPsych

Preben Bo Mortensen ^13, 29,30^; Ole Mors ^13,31^; Thomas Werge1^32^, 2008; Merete Nordentoft ^13,33^; David M. Hougaard^13,34^; Jonas Bybjerg-Grauholm^13,34^; Marie Bækvad-Hansen^13,34^;

13 iPSYCH, The Lundbeck Foundation Initiative for Integrative Psychiatric Research, Denmark

29 National Centre for Register-Based Research, Aarhus University, Aarhus, Denmark

30 Centre for Integrated Register-based Research, Aarhus University, Aarhus, Denmark

31 Psychosis Research Unit, Aarhus University Hospital, Aarhus, Denmark

32 Institute of Biological Psychiatry, MHC Sct. Hans, Mental Health Services Copenhagen, Roskilde, Denmark

33 Mental Health Services in the Capital Region of Denmark, Mental Health Center Copenhagen, University of Copenhagen, Copenhagen, Denmark

34 Center for Neonatal Screening, Department for Congenital Disorders, Statens Serum Institut, Copenhagen, Denmark

## 23andMe Research Team

Michelle Agee^3^, Stella Aslibekyan^3^, Robert K. Bell^3^, Katarzyna Bryc^3^, Sarah K. Clark^3^, Sarah L. Elson^3^, Kipper Fletez-Brant^3^, Pierre Fontanillas^3^, Nicholas A. Furlotte^3^, Pooja M. Gandhi^3^, Karl Heilbron^3^, Barry Hicks^3^, Karen E. Huber^3^, Ethan M. Jewett^3^, Yunxuan Jiang^3^, Aaron Kleinman^3^, Keng-Han Lin^3^, Nadia K. Litterman^3^, Marie K. Luff^3^, Matthew H. McIntyre^3^, Kimberly F. McManus^3^, Joanna L. Mountain^3^, Sahar V. Mozaffari^3^, Elizabeth S. Noblin^3^, Carrie A.M. Northover^3^, Jared O’Connell^3^, Aaron A. Petrakovitz^3^, Steven J. Pitts^3^, G. David Poznik^3^, J. Fah Sathirapongsasuti^3^, Janie F. Shelton^3^, Suyash Shringarpure^3^, Chao Tian^3^, Joyce Y. Tung^3^, Robert J. Tunney^3^, Vladimir Vacic^3^, Xin Wang^3^, Amir Zare^3^.

^3^23andMe, Inc. Sunnyvale, California, USA

## Extended Data

**Extended Data Fig. 1.**
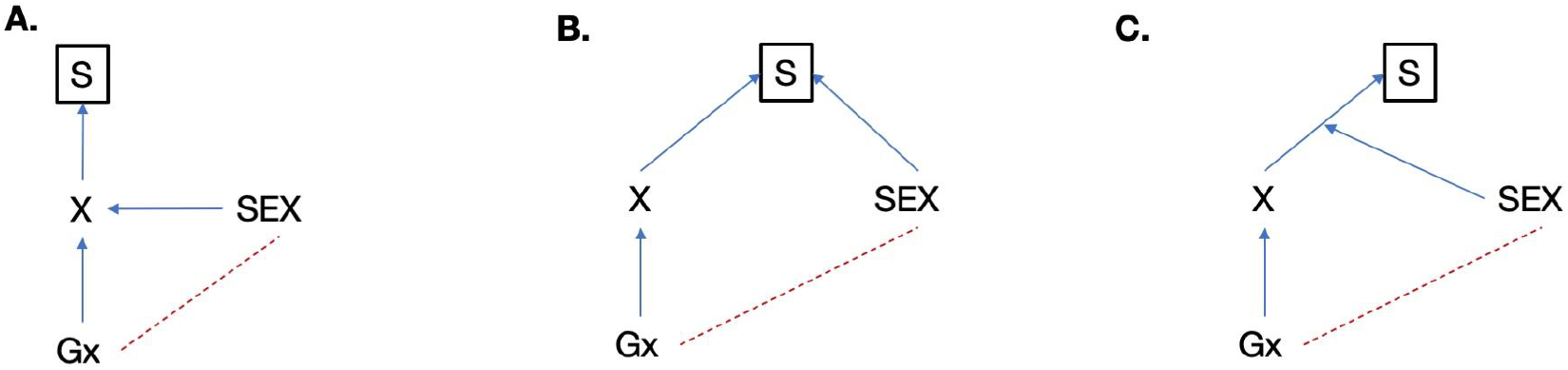
Different participation bias scenarios that may lead to a correlation between sex and genetic variants. S, selection (i.e. participation in the study); X, trait; Gx, genotype causing X. The assumed causal paths are shown in blue, and the induced correlations are shown in red. Three scenarios exist in which sex can become heritable due to selection. a, Sex causes X which in turn causes selection. b, X and sex influence the selection independently. c, The effect of X on selection is different between the two sexes. We have run simulations (Supplementary Fig. 3) and scenarios a and b are less likely to be observed because the effect of the trait on selection would need to be extremely large.

**Extended Data Fig. 2.**
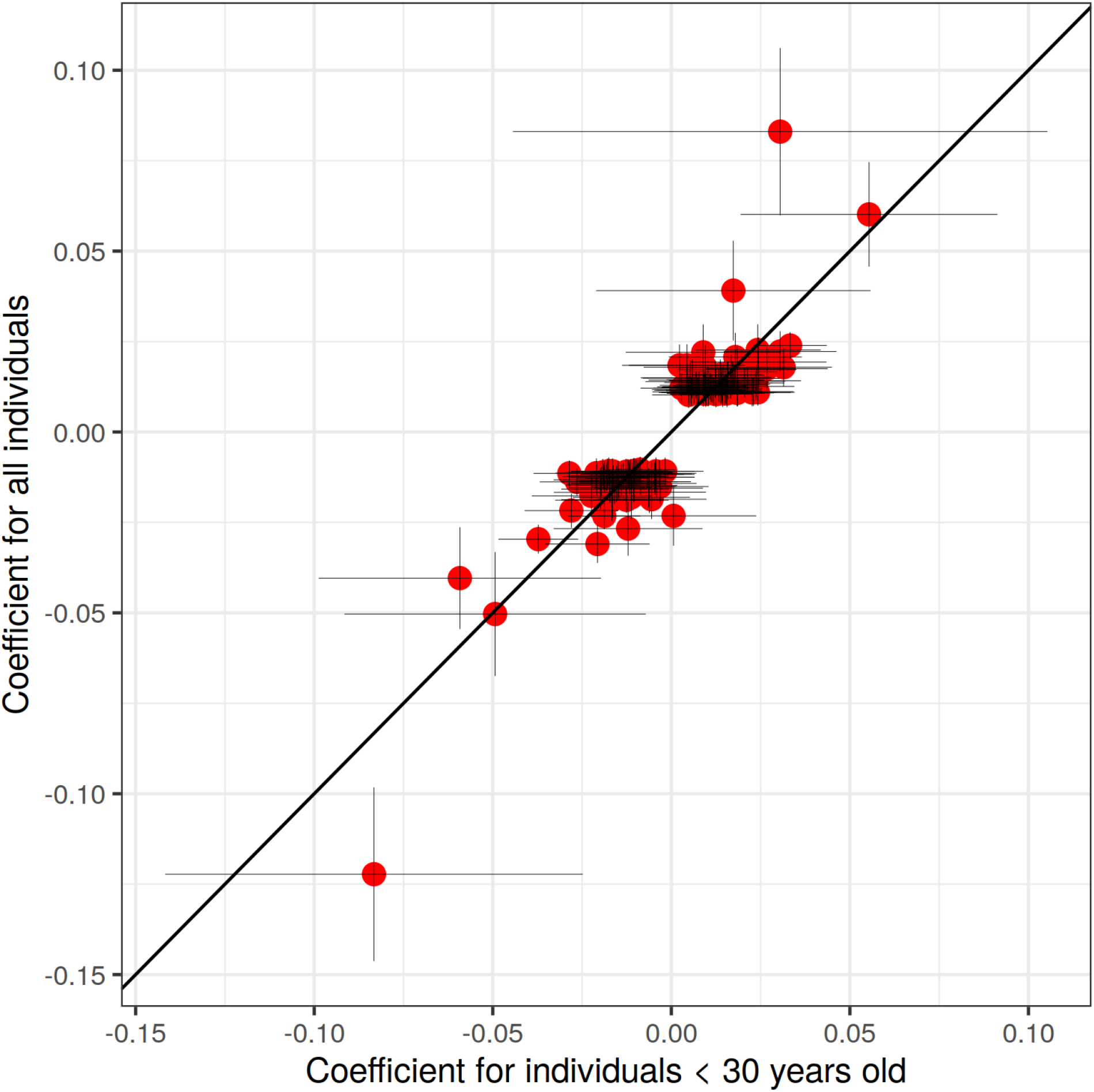
Effect size for association between SNPs and sex in 23andMe. On the y-axis is the effect in the entire study population (n = 2,462,132), and on the x-axis is the effect only among those younger than 30 years (n = 320,366). Error bars represent the confidence intervals for the effect size estimates.

**Extended Data Fig. 3.**
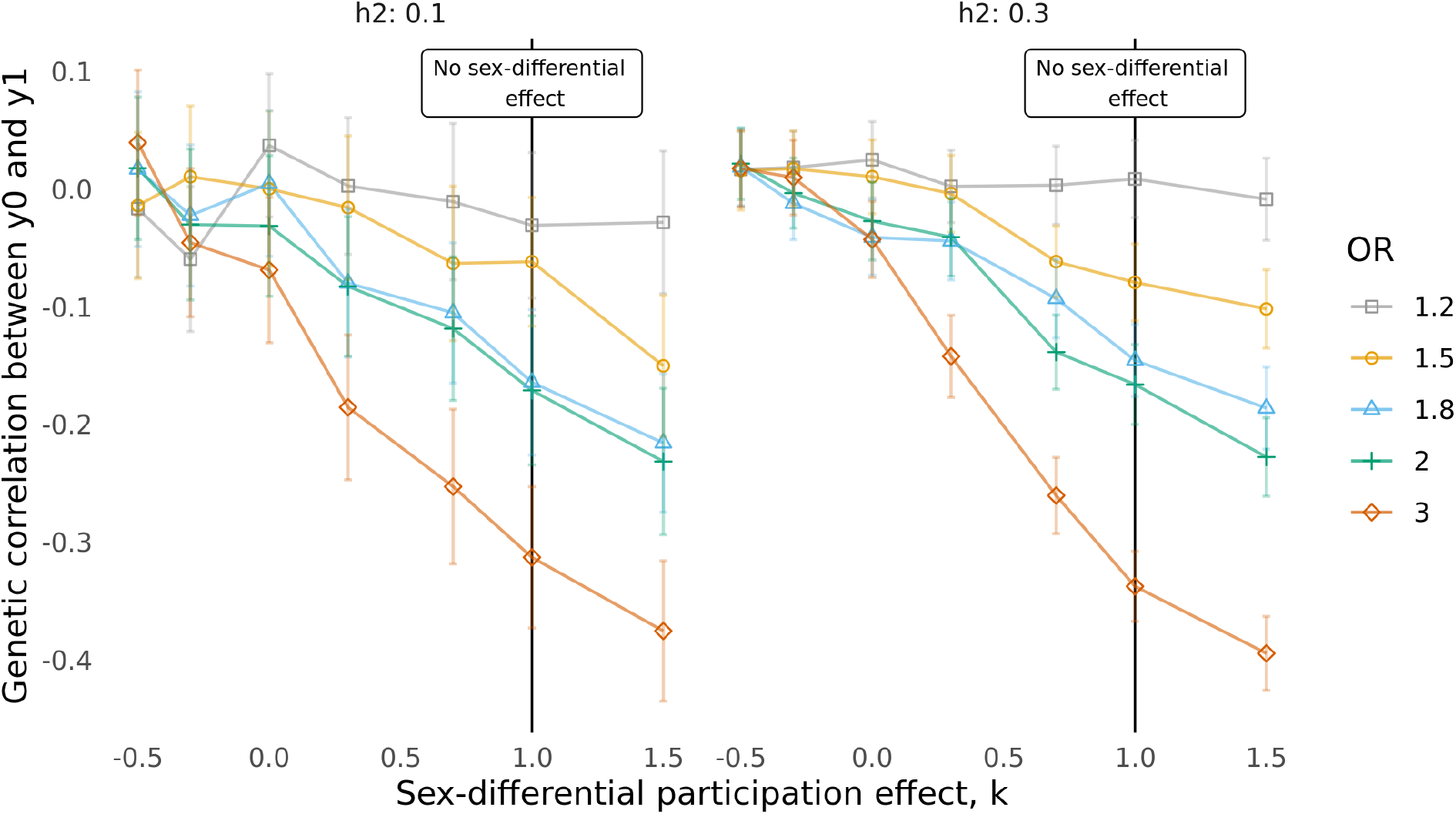
Effect of sex-differential participation bias on the genetic correlation between y0 and y1 when the phenotypes have h2 = 0.1 or h2 = 0.3. Each line represents a different degree of participation bias, expressed as the odds ratio (OR) used for the sampling. The higher the OR, the higher the degree of participation bias. The x-axis represents different values for the parameter k that gives the sex-differential effect. The smaller k is, the higher is the degree of the sex-differential effect.

**Extended Data Fig. 4.**
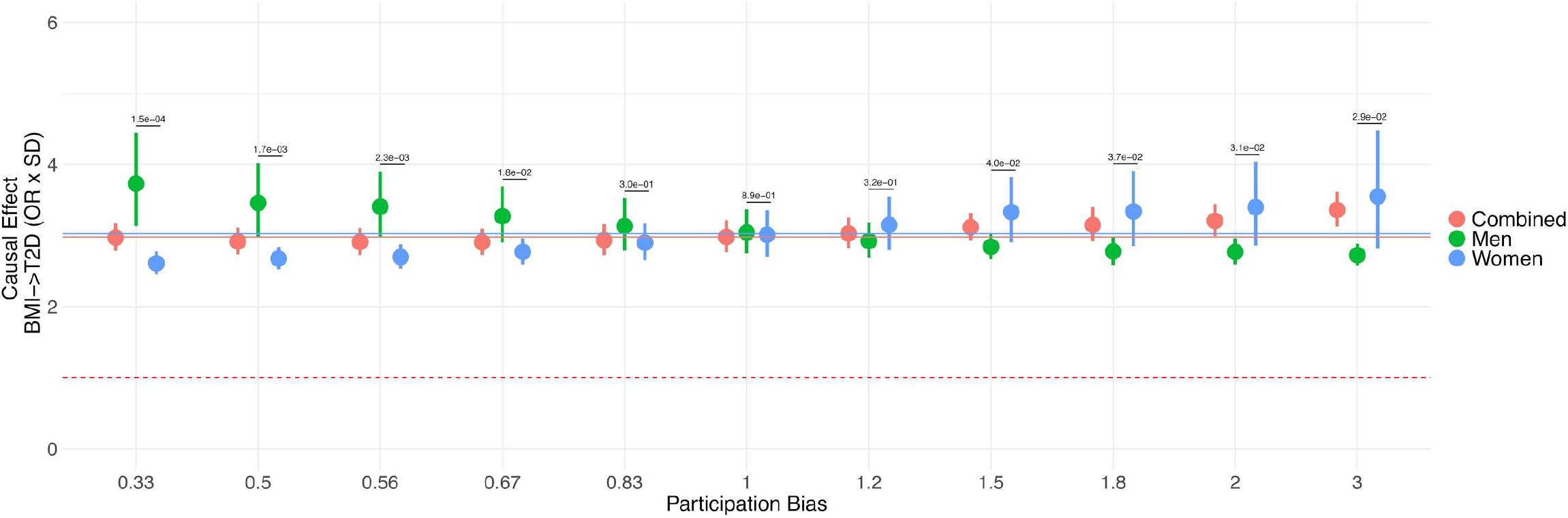
Effects of sex differential bias on the BMI →T2D relationship. The forest plot shows the effect of sampling men and women differentially based on BMI. The x-axis represents different values of bias introduced. For higher values, heavier males and leaner women are randomly picked. The number on top of the segment represents the P-value of the difference in effect between the two sexes using the Z-score method. The bias becomes large enough to be detected as “significant” even at the lower values of bias applied. The straight lines represent the effect of BMI on T2D estimated without any sample selection.

### Contributing cohorts

#### 23andMe

##### Cohort description

23andMe Inc. is a personal genetics company founded in 2006. Data for this study were available for approximately 2,462,000 individuals of European ancestry who provided informed consent and answered surveys online according to a human subjects protocol approved by Ethical & Independent Review Services, a private institutional review board. In this study we included 1,301,549 females and 1,160,583 males. Male (coded ‘0’) or female (coded ‘1’) case-control status was defined based on the concordance between the sex chromosomes and self reported sex. For configurations other than XX and XY, X0 was classified as female and XYY, XXY as male.

##### Genotyping and imputation

###### Genotyping

Genotyping was performed on various genotyping platforms: V1 and V2 Illumina HumanHap550+Beadchip (560,000 markers), V3 Illumina OmniExpress+Beadchip (950,000 markers), V4 custom (570,000 markers) and V5 Illumina Infinium Global Screening Array (~640,000 SNPs) supplemented with ~50,000 SNPs of custom content.

###### Imputation

We combined the May 2015 release of the 1000 Genomes Phase 3 haplotypes^1^ with the UK10K imputation reference panel^2^ to create a single unified imputation reference panel. To do this, multiallelic sites with N alternate alleles were split into N separate biallelic sites. We then removed any site whose minor allele appeared in only one sample. For each chromosome, we used Minimac3^3^ to impute the reference panels against each other, reporting the best-guess genotype at each site. This gave us calls for all samples over a single unified set of variants. We then joined these together to get, for each chromosome, a single file with phased calls at every site for 6,285 samples. Throughout, we treated structural variants and small indels in the same way as SNPs.

In preparation for imputation, we split each chromosome of the reference panel into chunks of no more than 300,000 variants, with overlaps of 10,000 variants on each side. We used a single batch of 10,000 individuals to estimate Minimac3 imputation model parameters for each chunk.

To generate phased participant data for the v1 to v4 platforms, we used an internally-developed tool, Finch, which implements the Beagle graph-based haplotype phasing algorithm^4^, modified to separate the haplotype graph construction and phasing steps. Finch extends the Beagle model to accommodate genotyping error and recombination, in order to handle cases where there are no consistent paths through the haplotype graph for the individual being phased. We constructed haplotype graphs for all participants from a representative sample of genotyped individuals, and then performed out-of-sample phasing of all genotyped individuals against the appropriate graph.

### UK Biobank

#### Cohort description

The UK Biobank cohort is a population-based cohort of approximately 500,000 participants that were recruited in the United Kingdom between 2006 and 2010^5^. Invitations to participate were sent out to approximately 9.2 million individuals aged between 40 and 69 who lived within 25 miles of one of the 22 assessment centers in England, Wales, and Scotland. The participation rate for the baseline assessment was about 5.5%. From these participants, extensive questionnaire data, physical measurements, and biological samples were collected at one of the assessment centers. In this study, we included 245,351 females and 206,951 males.

#### Genotyping and imputation

We used genotype data from the May 2017 release of imputed genetic data from the UK Biobank. The quality control and imputation were performed by UK Biobank and have been described elsewhere^5^. Briefly, genotyped variants were filtered based on batch effects, plate effects, departures from HWE, genotype platform, and discordance across control replicates. Participant samples were excluded based on missing rate, inconsistencies in reported versus genetic sex, and heterozygosity based on a set of 605,876 high-quality autosomal markers. Imputation was performed using IMPUTE4 with the HRC UK10K and 1000 Genomes Phase 3 dataset used as the reference set. Male (coded ‘0’) or female (coded ‘1’) case-control status was defined based on the concordance between the sex chromomes and self reported sex.

### iPSYCH

#### Cohort description

The iPSYCH study is a population-based case-cohort sample extracted from a baseline cohort consisting of all children born in Denmark between May 1st, 1981 and December 31st, 2005^6^. Those eligible were singletons born to a known mother and resident in Denmark on their one-year birthday. Cases were identified from the Danish Psychiatric Central Research Register (DPCRR)^7^, which includes data on all individuals treated in Denmark at psychiatric hospitals (from 1969 onwards) as well as at outpatient psychiatric clinics (from 1995 onwards). Cases were identified with schizophrenia, bipolar affective disorder, affective disorder, ASD and ADHD up until 2012. The controls constitute a random sample from the set of eligible subjects. The average (standard deviation) age of the individuals at recruitment (1st January 2012) was 18.3 (6.38) for males and 20.5 (6.16) for females. In this study, we included 31,012 females and 34,879 males.

#### Genotyping and imputation

##### Genotyping

Genotyping was performed at the Broad Institute (Cambridge, MA, USA) using the PsychChip array from Illumina (CA, San Diego, USA) according to the instructions of the manufacturer. Genotyping was carried out on the full iPSYCH sample in 23 waves and so was the subsequent data processing. Genotype calling of markers is described elsewhere (https://sites.google.com/a/broadinstitute.org/ricopili/utilities/merge-calling-algorithms). Prior to the subsequent QC and imputation SNPs were excluded when they were on either of two lists: a) a global blacklist comprising SNPs for which genotyping failed in 4 cohorts genotyped at the Broad as part of the PsychChip project (Psychiatric Genomics Consortium) with Illumina’s PsychChip and/or b) a local blacklist of SNPs for which the MAF in the GenCall and Birdseed call sets where substantially different (ΔMAF > 5%) prior to the merging of variants.

##### Imputation

Before subsequent imputation, the data was (strand) aligned with the respective reference sample. Phasing was achieved using SHAPEIT v2^8^ and imputation was done by IMPUTE2^9^ with haplotypes from the 1000 Genomes Project, phase 3 (1kGP3) as reference.

### Finngen

#### Cohort description

FinnGen is a public-private partnership project combining genotype data from Finnish biobanks and digital health record data from Finnish health registries (https://www.finngen.fi/en). Six regional and three country-wide Finnish biobanks participate in FinnGen. Finngen also includes data from previously established populations and disease-based cohorts. However, since we are interested in “passive” participation, we excluded individuals enrolled via epidemiological studies and only considered “passive”, hospital-based recruitments. We used genotype and phenotype data of 150,831 participants (86,694 females and 64,137 males), excluding population outliers via PCA. Finngen participants ages ranged from 18 to 110 years.

#### Genotyping and imputation

##### Genotyping

Samples were genotyped with Illumina (Illumina Inc., San Diego, CA, USA) and Affymetrix arrays (Thermo Fisher Scientific, Santa Clara, CA, USA). Genotype calls were made with GenCall and zCall algorithms for Illumina and AxiomGT1 algorithm for Affymetrix data. Genotyping data produced with previous chip platforms and reference genome builds were lifted over to build version 38 (GRCh38/hg38) following the protocol described here: dx.doi.org/10.17504/protocols.io.nqtddwn. In sample-wise quality control, individuals with ambiguous sex, high genotype missingness (>5%), excess heterozygosity (+-4SD) and non-Finnish ancestry were removed. In variant-wise quality control variants with high missingness (>2%), low HWE P-value (<1e-6) and minor allele count, MAC<3 were removed. Chip genotyped samples were pre-phased with Eagle 2.3.5 (https://data.broadinstitute.org/alkesgroup/Eagle/) with the default parameters, except the number of conditioning haplotypes was set to 20,000.

##### Genotype imputation with a population-specific reference panel

High-coverage (25-30x) WGS data (N= 3,775) were generated at the Broad Institute and at the McDonnell Genome Institute at Washington University; and jointly processed at the Broad Institute. Variant call sets were produced using the GATK HaplotypeCaller algorithm by following GATK best-practices for variant calling. Genotype-, sample- and variant-wise QC was applied in an iterative manner by using the Hail framework (https://github.com/hail-is/hail) v0.1 and the resulting high-quality WGS data for 3,775 individuals were phased with Eagle 2.3.5 as described above. Genotype imputation was carried out by using the population-specific SISu v3 imputation reference panel with Beagle 4.1 (version 08Jun17.d8b, https://faculty.washington.edu/browning/beagle/b4_1.html) as described in the following protocol: dx.doi.org/10.17504/protocols.io.nmndc5e. Post-imputation quality-control involved non-reference concordance analyses, checking expected conformity of the imputation INFO-values distribution, MAF differences between the target dataset and the imputation reference panel and checking chromosomal continuity of the imputed genotype calls.

### Biobank Japan

#### Cohort description

The BioBank Japan Project (https://biobankjp.org/english/index.html) is a national hospital-based biobank started in 2003 as a leading project of the Ministry of Education, Culture, Sports, Science and Technology, Japan. The BBJ collected DNA, serum and clinical information from approximately 200,000 patients with any of 47 target diseases between 2003 and 2007. Patients were recruited from 66 hospitals of 12 medical institutes throughout Japan (Osaka Medical Center for Cancer and Cardiovascular Diseases, the Cancer Institute Hospital of Japanese Foundation for Cancer Research, Juntendo University, Tokyo Metropolitan Geriatric Hospital, Nippon Medical School, Nihon University School of Medicine, Iwate Medical University, Tokushukai Hospitals, Shiga University of Medical Science, Fukujuji Hospital, National Hospital Organization Osaka National Hospital, and Iizuka Hospital). All patients were diagnosed by physicians at the cooperating hospitals. Details of study design, sample collection, and baseline clinical information were described elsewhere^10,11^.

#### Genotyping and Imputation

We genotyped samples using i) the Illumina HumanOmniExpressExome BeadChip or ii) a combination of the Illumina HumanOmniExpress and the HumanExome BeadChip. We applied standard quality-control criteria for samples and variants as described elsewhere^15^. The genotypes were prephased using Eagle and imputed using Minimac3 with a reference panel using a combination of the 1000 Genomes Project Phase 3 (version 5) samples (n = 2,504) and whole-genome sequencing data of Japanese individuals (n = 1,037)^12^.

## Participation bias simulations

To assess the effects of sex-differential participation bias we devised a sampling strategy to modulate the degree of bias and applied it to simulated data.

We used genotype data of 350,000 unrelated individuals of European ancestry from UKBB and 1,159,813 common HapMap variants to generate two synthetic phenotypes, y0 and y1. To ensure the phenotypes were uncorrelated with sex and had the same proportion of males and females, we first assigned to each individual a dummy variable representing sex, drawing values from a binomial distribution with p=0.5.

The phenotypes were simulated using the infinitesimal model^13^ as implemented in Hail version 0.2.24, which assumes that the genetic component of a trait comes from a large number of small effects:

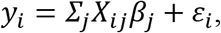

where *y_i_* is the phenotype of individual *i, X_ij_* is the genotype of individual *i* at SNP *j, β_j_* is the effect size of SNP *j* and *ε_i_* is environmental noise. SNP effect sizes are modelled as normally distributed with mean 0 and variance equal to the imposed SNP-heritability divided by the number of SNPs, *M*:

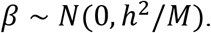

We looked at the effects of moderate and higher heritability, with values of *h*^2^ =0.1 and *h*^2^ =0.3 for both traits. In both cases the traits were simulated as genetically uncorrelated.

### Sampling strategy

We aimed at simulating the effects for all 3 models reported in figure S1.

### Participation bias on a trait which shows differences between males and females (Model A)

We verified if the observed effects could be generated by simple selection bias on a trait which shows differences between males and females (Extended Figure 1 model A). We simulated a trait *X* with different heritability values of 0.1, 0.3 and 0.8 as described above. We then added an effect of 0.5 and 1 standard deviations in one sex as follows:

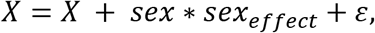

where *ε* is random normally-distributed noise, *ε* ~ *N*(0,0.01). We then sampled the population as described above but applying selection only on one trait (*X*) and without sex-differential effects:

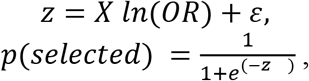

with OR=[1.2, 1.5, 2, 3, 5].

For each subsample we estimated the heritability for sex on the liability scale and reported the results in Supplementary Figure 4.

Additionally, we compared the educational attainment in the US census and in 23andMe, defined as years of education as follows:

Less than high school: 10
High school: 12
Associate/vocational/some college: 14
Bachelor: 16
Master/professional: 19
Doctorate: 22

### Participation bias is influenced by a trait and sex independently (Model B)

We verified if the observed effects could be generated by selection bias both on a trait *X* and sex independently (Extended Figure 1 model B). We simulated a trait *X* with different heritability values of 0.1, 0.3 and 0.8 as described above. We then used the following model for selection:

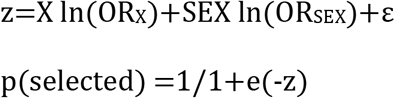

With OR_X_=2, OR_SEX_=[1.2,1.5,2,3,5] and ε~N(0, 0.01).

For each subsample we estimated the heritability for sex on the liability scale and reported the results in Supplementary Figure 2C.

### Participation bias is influenced by a trait in a sex specific manner (Model C)

**Supplementary Figure 1** shows the basic workflow to simulate the phenotypes y0 and *y1* and induce sex-differential participation bias. y0 and y1 are simulated to be genetically uncorrelated in the full population. Each individual is then assigned to a probability of being selected as follows:

1. A variable *z* is computed as the weighted sum of the phenotypes:

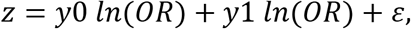

where *ε* is random normally-distributed noise, *ε* ~ *N*(0,0.01), and the odd ratio (OR) represents the degree of participation bias. The higher the OR, the higher the participation bias since more individuals with greater values of the phenotypes will be selected. OR=1 represents the case when no participation is induced.
2. A sex-specific effect is given multiplying *z* by the parameter *K* in one sex:

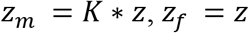 Lower (negative) values of *K* represent an higher sex-differential bias. *K*=0 and *K*=1 represent two special cases where, respectively, one sex is sampled randomly and both sex are sampled equally (no sex-specific bias).
3. The probability associated to each individual is computed as the logistic function of the sex-specific *z*

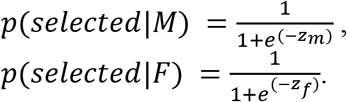

We used different combinations of the parameters *K* ([-0.5,-0.3,0,0.3,0.7,1,1.5]) and *OR* ([1.2, 1.5, 1.8, 2, 3]) to control the degree of bias induced. At each step the subsampled population contained nearly half of the original population.

Results for all 3 scenarios are reported in **Supplementary Figure 2**

### Results

#### Heritability of sex and Mendelian Randomization

**Figure 3B** shows how sex becomes heritable at the increasing of participation bias (keeping sex-differential effect fixed.). Moreover, a causal effect between y0 and sex is induced, as shown is **Figure 3C** for OR=1.8.

We reported the complete results from the simulations in **Supplementary Table 9.** As expected, with the increasing of participation bias also the SNP-heritability for sex increases and becomes significant.

#### Genetic correlation between y0 and y1

**Extended Figure 3** shows that a spurious negative genetic correlation between the traits y0 and y1 (simulated as genetically uncorrelated) is induced and this effect increases at the increase of both parameters. Moreover, as shown in **Supplementary Figure 3**, this effect is exacerbated when adjusting for sex. However, this is issue arises only when there is a substantial sex-differential effect and in a realistic scenario (see *Consistency between our simulation parameters and real data*) corresponding to OR=1.2 and k=0.7 none of the mentioned effects is observed.

#### Genetic correlation between males and females for a given phenotype

**Supplementary Figure 5** reports the genetic correlation between males and females for y0 and y1. This shows how participation bias does not arise any effect when stratifying the analysis for sex.

#### Consistency between our simulation parameters and real data

Our simulation strategy was designed to provide realistic scenarios of sampling bias. We used the differences in educational attainment (EA) between those UK Biobank individuals that participated in all the online 24-h diet follow up questionnaires *vs*. those that did not (**Supplementary Table 8)** and compared it with the differences in sampled and non-sampled individuals for y0 and y1 obtained from simulations. However, results reported in **Supplementary Table 8** are on the observed scale and therefore not directly comparable with results from simulations, which use standardized variables. Thus, we standardized EA and obtained standardized differences between individuals that participated in all the online 24-h diet follow up questionnaires *vs*. those that did not of 0.30 and 0.37 standard deviations, in males and females respectively. Next, we assessed which OR and *k* parameter in the simulations would provide similar changes between the original group and the “sampled” group. In our simulation, the closest value to these differences was observed for an OR=1.25 and a *k* parameter=0.7.

## Sex-specific MR analysis

In order to verify the impact of sex differential participation bias on causal inference through MR using real data, we imposed additional bias to a real example from the literature^14^. We focused on the sex-specific causal relationship between body mass index (BMI) and Type 2 diabetes (T2D) recently reported by Censin and colleagues^35^. In the original paper, the authors report a strong difference in the effect of BMI on T2D in men and women (p=1.4×10^−5^). We thus wondered if this could be explained by sex differential selection on BMI. That is, if changing the degree of sex-differential selection on BMI would change the sex-specific estimates.

We first notice that Cansin and colleagues standardized BMI separately for males and females. They thus found a larger odds ratio (OR) for T2D per standardized increase in BMI genetic score in females (3.77) than in males (2.79). However, the standard deviation of BMI in UK Biobank is larger in females (~5.1 kg/m^2^) than in males (~4.2 kg/m^2^), and we find that this sex difference in the variance of BMI accounts for the apparent sex difference in the effect of BMI on T2D risk. In an alternative approach, using exactly the same UK Biobank data, we scaled the BMI in males and females to the same sex-combined standard deviation (~4.75 kg/m^2^) and observed no difference in the effect of BMI genetic score on T2D risk between males and females (OR 3.03 *vs* 3.03). Therefore BMI contributes to a smaller proportion of the liability to T2D in women than it does to men as it has a wider phenotypic distribution, but importantly a one unit increase of BMI is not different in terms of OR for T2D risk in women than men. From this difference in interpretation lies the difference we observe in the results. Nonetheless, the goal of this analysis is to show that a sex-specific causal effect can be induced by sex-differential participation bias. Thus, bias was introduced differently for men and women based on the standardized BMI.

We used the same sampling strategy described in **Participation bias simulations** and with *K*=([−0.5, −0.3,0,0.3,0.7,1,1.5]) and OR=([0.33,0.5,0.56,0.67,0.83,1,1.2,1.5,1.8,2,3]). These OR values are symmetric around 0 on the *ln(OR)* scale. This sampling choice was selected because of the different prevalence between men and women at baseline.

In order for our results to be comparable with the published results MR estimates were obtained using the Wald ratio method while SE were estimated using the delta method and second order weights. The Wald ratio method consists of running the regression of the exposure trait on it’s instrument (the polygenic score (PGS) in our case) and then the logistic regression between the outcome and the instrument.

The causal estimate is then estimated by the ratio of 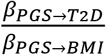

For each regression, we used as covariates array type, batch of genotyping, 40 Principal components, age, age^2^, and sex only for the combined analysis.

The overall and sex-specific weights were obtained from the supplementary material in the Censin and colleagues paper^35^. As an outcome, we used T2D using “probable” and “possible” cases as defined in the algorithm from Eastwood and colleagues^15^. **Extended Figure 4** reports the results for K=-1 while full results can be found in **Supplementary table 10**

## GWAS of 565 heritable traits with and without adjustment for sex

To determine trait heritability we used the approach by Walters and colleagues^16^. In particular we select 565 traits with *confidence=“high”* and *significance level ≥ “z4”* and available in both sexes. Individuals included in the analyses and methods used to run the GWASs are described at http://www.nealelab.is/uk-biobank/ukbround2announcement, with the only difference that the following covariates were used: 20 principal components and age. We ran two set of GWASs: one including sex as a covariate, the other without including sex as a covariate. We conducted two main analyses on these results; First, for the same trait, we calculate genetic correlation between the GWAS adjusted and non-adjusted by sex. Second, we calculate the genetic correlations between each trait and all the other traits for the two sets of GWASs (adjusted and non-adjusted by sex) using a faster version of LDscore regression v. 1.0.0 (https://github.com/astheeggeggs/ldsc). We then compared the two correlation profiles.

## Proposed correction methods and implementation in GenomicSEM

### Heckman correction generalization

Participation bias is a known problem in epidemiology and econometrics and several corrections have been proposed. Heckman correction^17^ is widely used in econometrics, but only recently re-discovered in epidemiological research^18,19^.

We are interested in the relationship between Y (outcome) and X (exposure) but we only observe these variables among individuals that participated in the study (S=1). If the variables were observed only among study participants, we use the notation Y* and X*.

The main challenge is that the distribution of Y* in the entire population is not available.

Heckman correction addresses this challenge in two steps.

First, fit a probit model of participation:

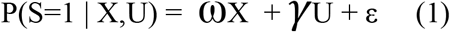

Where the probability of participating S=1 depends on some explanatory variables, X the variable of interest and other variables U related to S but independent of Y. It is important that the model includes at least one U variable to act as an instrumental variable (i.e. another phenotype not related to the outcome of interest if not through participation) and avoid excessive collinearity with X.

For each participant an expected probability of participation P(S) is then obtained based on (1).

Second, the expected probability of participation among individuals selected in the study P(S*) is used as covariates in the model when testing the association between X* and Y*. Given that we have assumed a probit model, the resulting distribution will be a truncated normal we first estimate the inverse mills ratio of the predicted probabilities only in the selected samples.

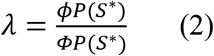

Where φ denotes the standard density function while Φ is the standard normal cumulative distribution function.

λ is then added in the regression as covariate:

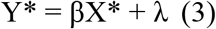

The problem can be at this point simplified by retrieving the correlation matrix between Y*, X* and S*.

Given that under the “Cheverud’s Conjecture”^20,21^ genetic correlations can be used as proxies of phenotypic correlations, we can use the genetic correlations obtained from LDscore regression for the GWAS of Y*, X* and λ to fit (3) using GenomicSEM^22^.

However, there are several limitations. First, we need to assume an underlying bias model which should include at least X and an instrument variable U. The latter might be challenging to obtain. Second, X and any additional variable used to calculate P(S) have to be measured in the population of interest and the analysis must be limited to the samples which have all these variables. These two conditions may not always be easy to achieve and thus a method which does not require them is more desirable.

### Genomic structural equation model to estimate the genetic correlation between pairs of traits despite the presence of collider bias induced by selection on both traits

As an illustration of the modeling possibilities afforded by access to the allele frequency those that do not participate in a study or biobank we construct a genomic structural equation model that corrects for collider bias induced by sample selection.

We are interested in the relationship between Y (outcome) and X (exposure) but we only observe these variables among individuals that participated in the study (S=1). If the variables were observed only among study participants, we use the notation Y* and X*.

If the probability of selection into the sample is caused by X and Y, then selection results in collider bias of the relationship between Y* and X*. If the effects of Y and X on S are positive (e.g. higher X or Y results in selection into the sample) and both X and Y are heritable traits, a negative genetic correlation is induced between Y* and X*^23^.

Suppose we obtain the summary statistics for 3 GWAS: a GWAS of Y*, X* and a GWAS where the sample allele frequency is compared to the true population allele frequency (S).

We can then construct a 3*3 genetic covariance matrix of Y*, X* and S, where S positively correlates with Y* and X* and, due to collider bias, Y* and X* negatively correlate. Important to note is that the positive effects of Y and X on S are what cause the negative correlation between Y* and X* and are proportional to it, a bigger effect on S, or stronger selection, induce stronger collider bias and negative (genetic) correlation between Y* and X*.

We use GenomicSEM to fit a path model which only allows for a single path for the assortation between S and X, Sand Y and X and Y:

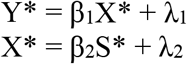

Where we constrain:

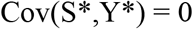

Because the estimate 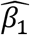 needs to accommodate the positive association between S* and Y*, and the (proportional) negative association between Y* and X* induced by collider bias, the cancels these quantities out and is ± equal to β_t_ in the regression:

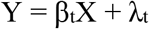

This result is validated in a simulation described below. The model assumes no unmeasured confounders distort the relationship between Y* and S or the relationship between X* and S. The model could be extended based on instrumental variable techniques to accommodate the presence of unmeasured confounders. However, as the model merely exists to illustrate the value of observing the true population allele frequencies extending the model for such eventualities is beyond the scope of the current paper. Code to fit the model is found here: https://github.com/dsgelab/genobias.

### Application to simulated data

To validate the two correction approaches we propose, we simulated phenotypes X and Y (h^2^=0.3) to have rg=[−0.3, −0.1, 0, 0.1, 0.3]. For each case we induced participation bias as described in **Participation bias simulation**, with OR=[1.2, 1.5, 1.8, 2, 3] and adding a trait U, also simulated with h2=0.3 but uncorrelated with both X and Y, to determine sample selection:

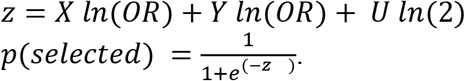

Simulations results are shown in **Supplementary Table 11**.

### Collaborators

#### iPsych

Preben Bo Mortensen 1, 2, 9, 10; Ole Mors 1, 6; Thomas Werge1, 7, 2008; Merete Nordentoft 1, 11; David M. Hougaard 1, 5; Jonas Bybjerg-Grauholm 1, 5; Marie Bækvad-Hansen 1, 5;

1 iPSYCH, The Lundbeck Foundation Initiative for Integrative Psychiatric Research, Denmark

5 Center for Neonatal Screening, Department for Congenital Disorders, Statens Serum Institut, Copenhagen, Denmark

6 Psychosis Research Unit, Aarhus University Hospital, Aarhus, Denmark

7 Institute of Biological Psychiatry, MHC Sct. Hans, Mental Health Services Copenhagen, Roskilde, Denmark

8 Department of Clinical Medicine, University of Copenhagen, Copenhagen, Denmark

9 National Centre for Register-Based Research, Aarhus University, Aarhus, Denmark

10 Centre for Integrated Register-based Research, Aarhus University, Aarhus, Denmark

11 Mental Health Services in the Capital Region of Denmark, Mental Health Center Copenhagen, University of Copenhagen, Copenhagen, Denmark

## Supplementary figures

**Supplementary Figure 1:**
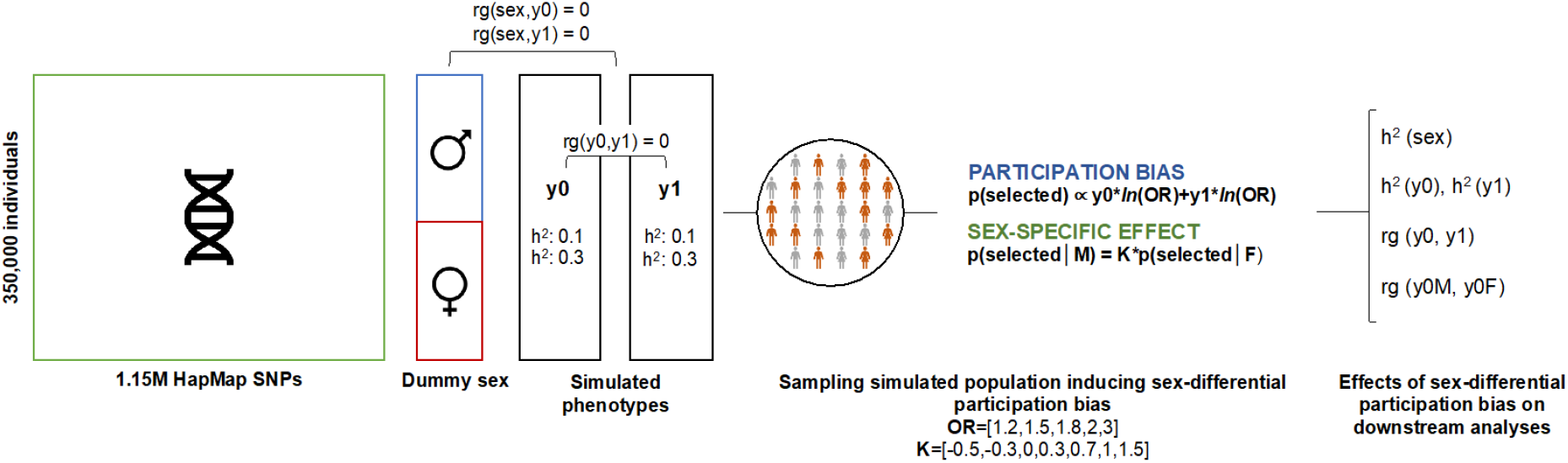
simulations pipeline. A dummy sex variable is assigned to 350,000 unrelated individuals from UKBB. Based on 1,159,813 HapMap variants, two genetically uncorrelated traits (*rg(yO,yl)=O*) with the same SNP-heritability are simulated. The simulated population is then sampled inducing sex-differential participation bias at different degrees, expressed in terms of the parameters OR (participation bias) and K (sex-specific effect). The consequences of sex-differential participation bias are assessed looking at the heritability of sex (*l·>2(sex)*) and of the simulated traits (*h2(yθ), h2(yl)*), the genetic correlation between the traits (*rg(yO,yl)*) and between males and females for a given trait (*rg(yOM,yOF)*).

**Supplementary Figure 2:**
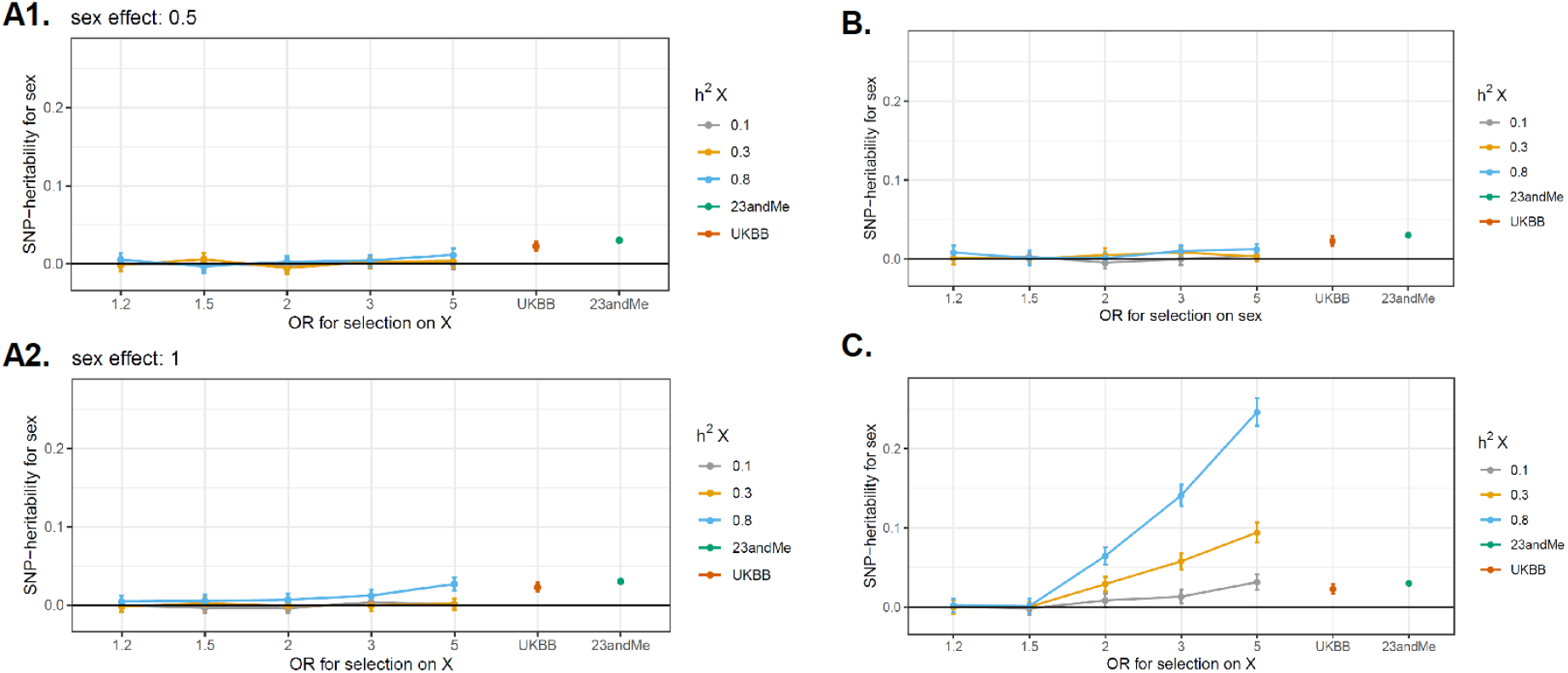
SNP-heritability estimates for sex (on liability scale) for the participation bias scenarios presented in Supplementary Figure 1. X is a trait simulated with heritability of 0.1,0.3 and 0.8 (N=350,000 simulated individuals). Varying the parameter OR, different degrees of selection are induced, accordingly to the specific scenario. In all the panels, the last two points on the X axis report the SNP-heritability for sex observed in UKBB and 23andMe. Error bars represent the confidence interval for the SNP-heritability estimate. **A1.** Sex causes X which in turn causes selection, with an effect of 0.5 standard deviation. **A2.** Sex causes X which in turn causes selection, with an effect of 1 standard deviation. **B.** X and sex influence the selection independently. C. Sex-differential participation bias: the effect of X on selection is different between the two sexes.

**Supplementary Figure 3:**
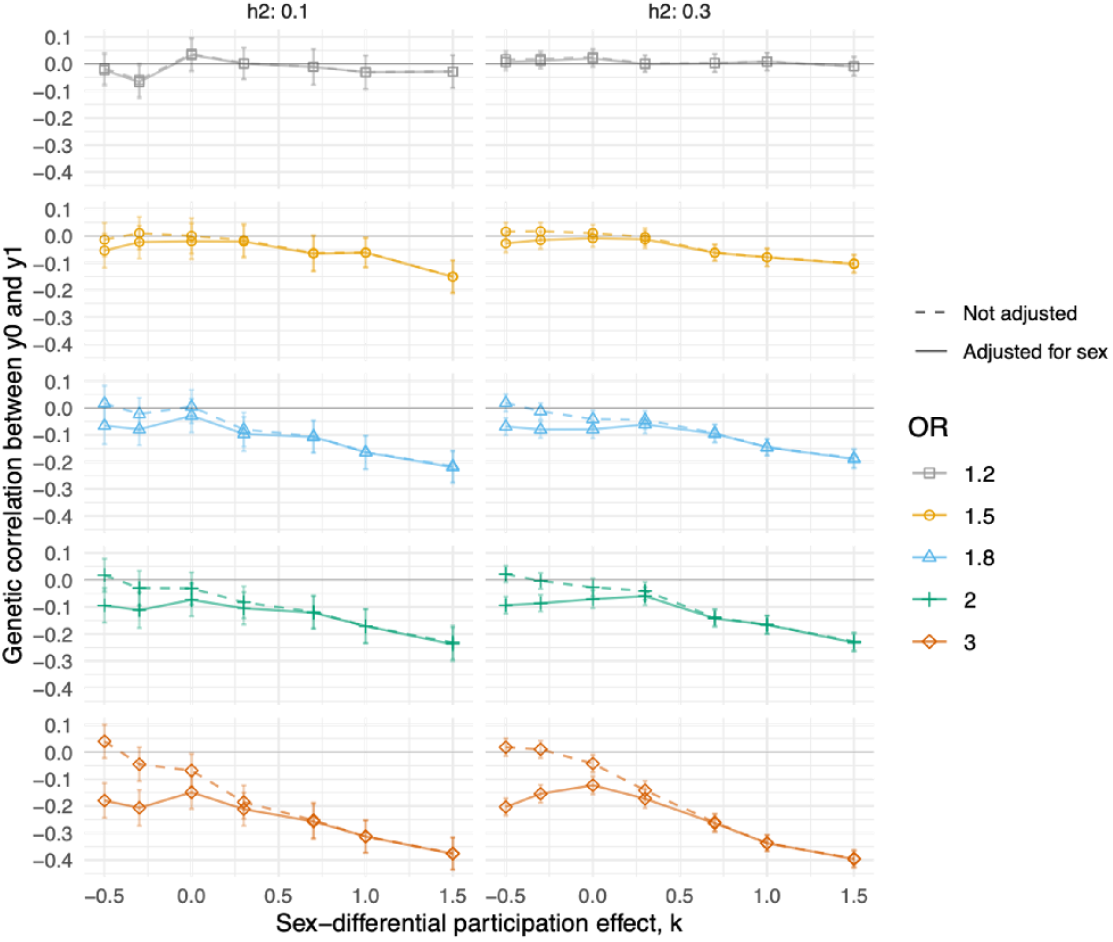
Effect of adjusting for sex when a spurious genetic correlation between y0 and yl is induced by participation bias, when the phenotypes have both h2=0.1 and h2=0.3. Each line represents a different degree of participation bias, expressed as the odd ratio (OR) used for the sampling. Higher the OR, higher the degree of participation bias. The x-axis represents different values for the parameter k, that gives the sex-differential effect. Adjusting for sex increases the degree of bias especially for lower values of k, for which the difference in the distribution of y0 and yl in the two sexes is greater.

**Supplementary Figure 4:**
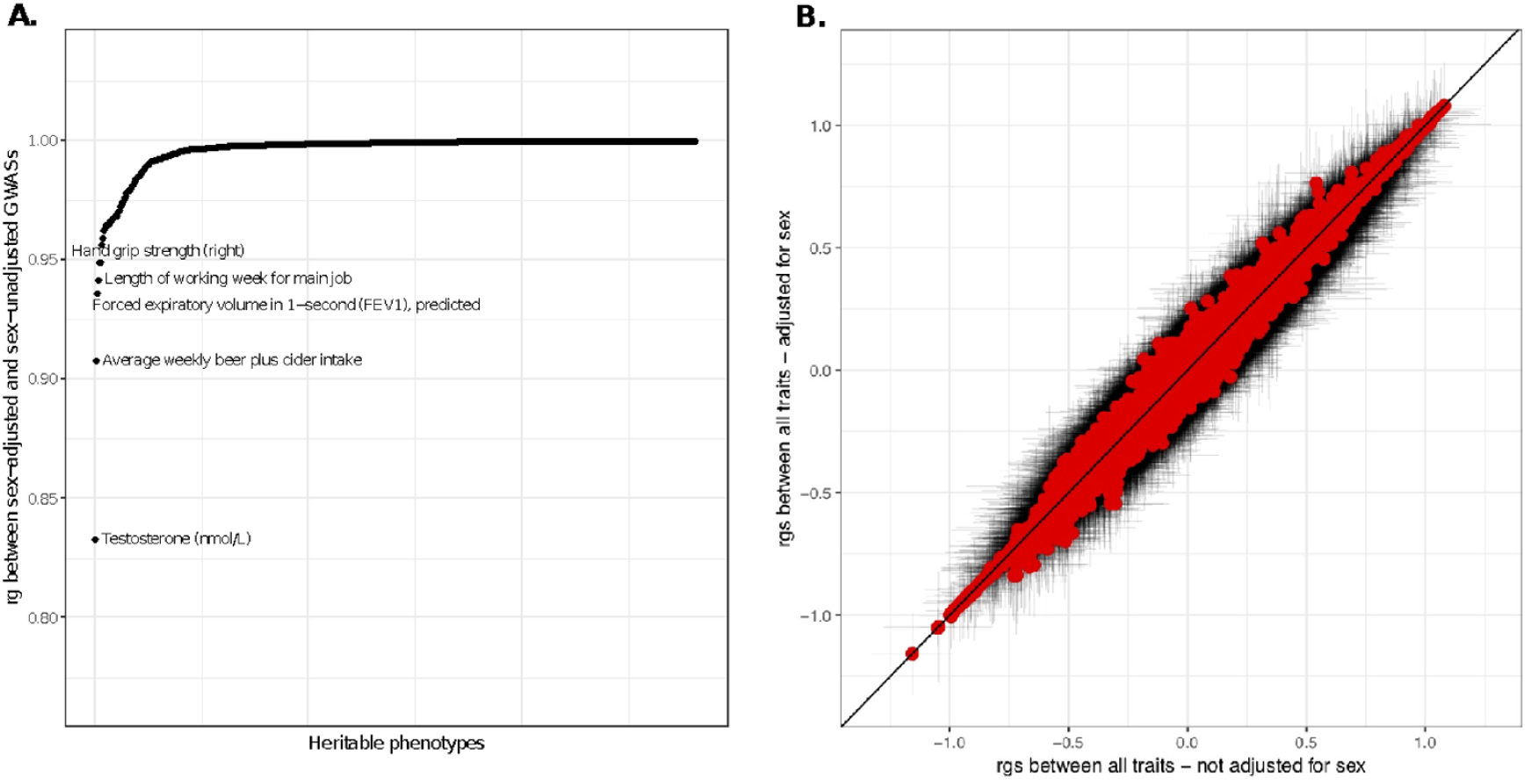
GWAS of 565 traits in UK Biobank with and without adjustment for sex. **Panel** A. genetic correlation between the sex-adjusted and sex-unadjusted GWAS. Top 5 traits with lowest genetic correlation are reported. **Panel B.** Genetic correlation between each trait and all the other, on the x-axis the results are from a sex unadjusted GWAS, on the y-axis the results are from a sex-adjusted GWAS. The horizontal and vertical bars represent confidence intervals.

**Supplementary Figure 5:**
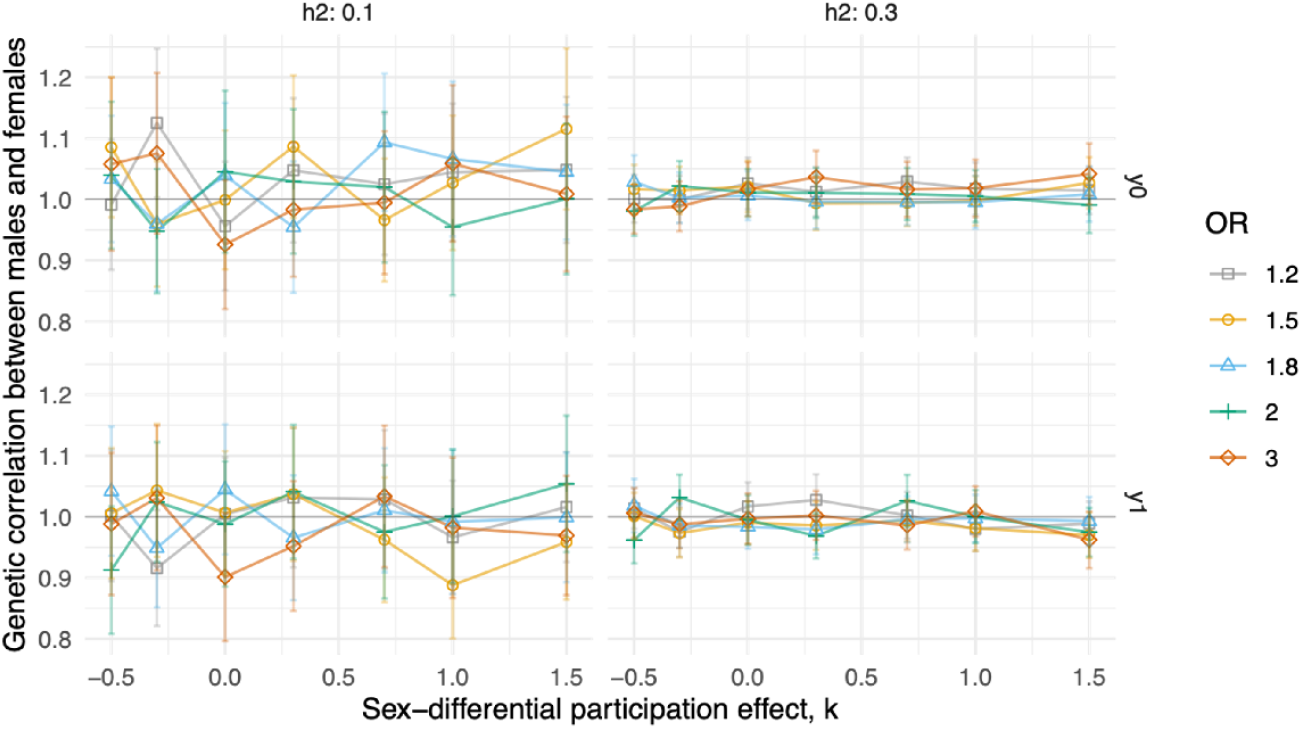
Genetic correlation between males and females for a given phenotype (N=350,000 simulated individuals). Each line represents a different degree of participation bias, expressed as the odd ratio (OR) used for the sampling. Higher the OR, higher the degree of participation bias. The x-axis represents different values for the parameter k, that gives the sex-differential effect. Sex-differential participation bias does not impact the genetic correlation between males and females. Error bars represent the confidence interval for the SNP-heritability estimate.

**Supplementary Figure 6:**
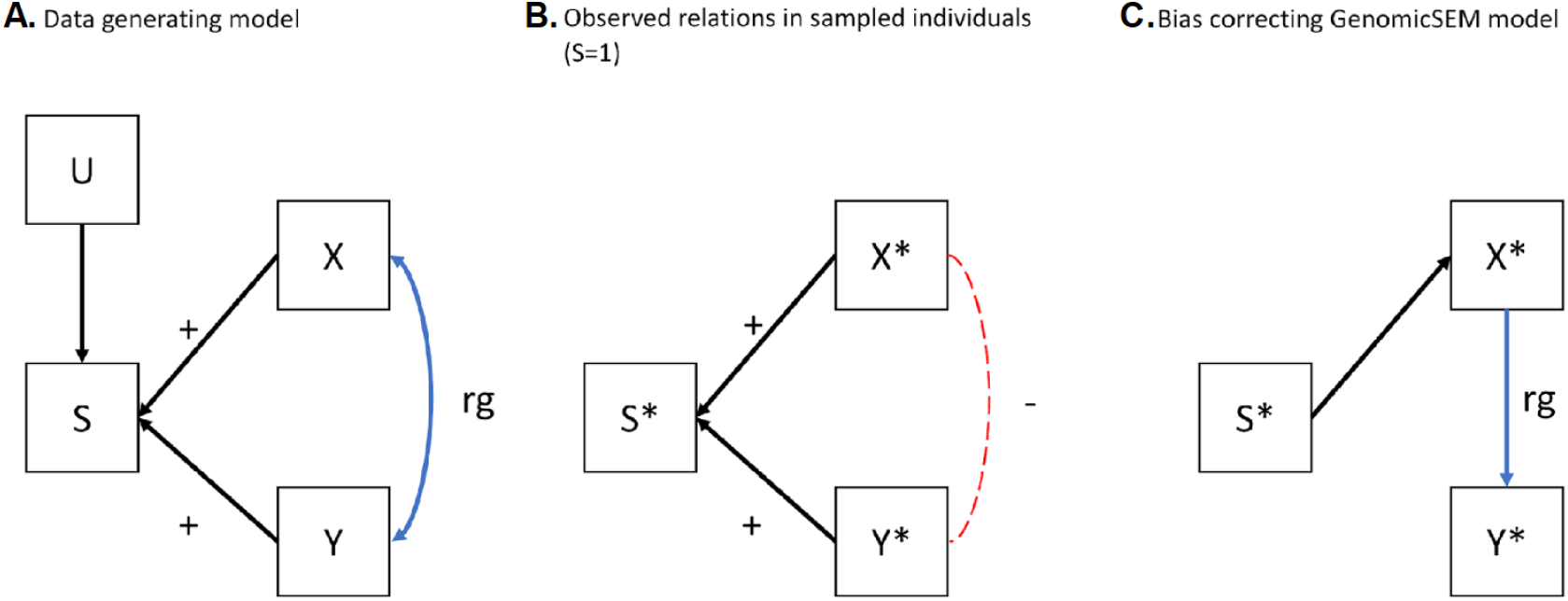
**A.** schematic representation of the data generating model, where S, the selection probability, is caused by X, Y and unmeasured variable(s) U. B. the expected relationships between GWAS of ×*,Y* and S*, where the * indicates the GWAS of Y and X are performed in selected individuals and the GWAS of S* is a GWAS of the dichotomous variable selected yes/no. C. the GenomicSEM model which forces the relationship between Y* and S*, as well as the relationship between X* and Y* through a single path, resulting in a corrected estimate of the relationship between X and Y based on Y*, X* and S*.

